# Realigning representational drift in mouse visual cortex by flexible brain-machine interfaces

**DOI:** 10.1101/2024.05.23.595627

**Authors:** Siyuan Zhao, Hao Shen, Shanshan Qin, Shouhao Jiang, Xin Tang, Madeleine Lee, Xinhe Zhang, Jaeyong Lee, Juntao Chen, Jia Liu

## Abstract

The ability to stably decode brain activity is crucial for brain-machine interfaces (BMIs), which are often compromised by recording instability due to immune responses and probe drifting. In addition, many brain regions undergo intrinsic dynamics such as “representational drift”, in which neural activities associated with stable sensation and action continually change over time. In this study, we employed tissue-like flexible electrode arrays for recording visual stimulus-dependent single-unit action potentials in the mouse visual cortex. The flexible electrode array enabled us to record action potentials from the same neurons over extended periods under visual stimuli, allowing us to characterize the representational drift during these stimuli. With this approach, we tested hypotheses about the origins and mechanisms of representational drift, tracked latent dynamics transformation, and modeled these dynamics with affine transformation. Our findings enabled the construction of a single, long-term stable, high-performance visual information decoder that accounts for representational drift, potentiating chronically stable flexible BMIs in brain regions experiencing representational drifts.

## Introduction

Brain-machine interfaces (BMIs) largely rely on stable brain activity decoding, which can be compromised by recording instability due to immune responses and probe drifting^1–3^. In addition, recent studies have indicated that many brain regions such as the posterior parietal cortex^4^ and hippocampus^5^ that are associated with decision making, learning, and memory, as well as sensory cortices such as olfactory cortex^6^ and visual cortex^7,8^ that encode external stimuli exhibit intrinsic dynamics such as “representational drift” in their neural activities. This phenomenon describes the variability in neural responses to consistent external variables such as behavioral output and sensory input. Consequently, recording instability compounded by intrinsic representational drift challenges the reliability of brain decoding. Traditional algorithms and calibration^9–11^ have been developed to compensate for probe instability and maintain long-term performance of BMIs, particularly for motor-related brain activity decoding. However, even stability of neural representations in motor-related brain regions over extended periods is not taken as ground truth based on several studies^12–14^.

To address the instability of recording, we recently developed ultra-flexible hexode mesh electrode arrays^15^. The mesh structure in ultra-flexible polymer ribbons can form an interwoven structure with soft brain tissue, enabling improved biocompatibility, reduced chronic immune response, and enhanced long-term recording stability. The hexode electrode array structure allows the identification and triangulation of the signals from the same neurons by multiple neighboring electrodes^6,9^. By leveraging the flexible hexode electrode array and data analysis pipeline, we achieved high-resolution, long-term stable tracking of the behavior-dependent electrical activities from the same neurons in the mouse primary visual cortex over multiple months.

These properties of our electrode arrays allow us to record neuron activities in the primary visual cortex, track the intrinsic changes in neurons’ visual response properties, and study representational drift over 5-6 months when mice were repetitively presented with the same natural movies (Fig. 1a-b). Our findings illustrate that the neural responses to natural movie stimuli show a persistent drift more explainable by the difference in stimulus experience numbers than time intervals between stimuli. Additionally, population-level latent dynamics in neural activities displayed long-term variability, which, however, can be approximated by a model of affine transformation over time (Fig. 1b). By stably tracking the neural dynamics from the same neurons and modeling the variability of latent dynamics with transformation, we were able to recover more consistent structures in neural activities underlying the same visual stimulus. As a result, we can compensate for the intrinsic change in the latent dynamics to achieve a high-performing information decoder in the visual cortex of the mouse brain. We envision this finding and application can be potentially translated to other brain regions known for their representational drift over extended periods.

**Fig. 1.**
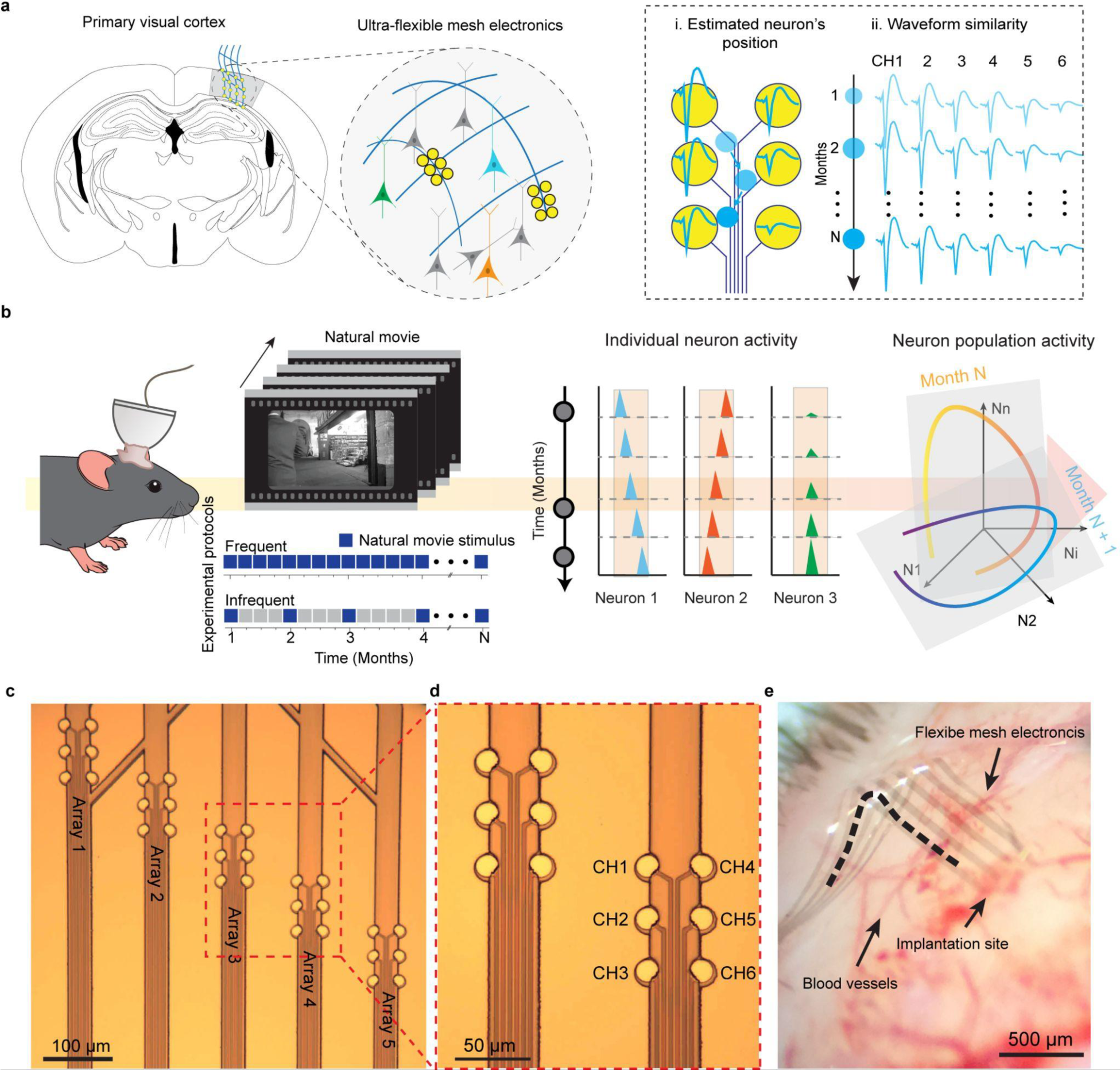
Long-term monitoring of representational drift in the mouse visual cortex using flexible electrode arrays. **a,** Schematics demonstrating the long-term stable tracking of neural activities from the same neurons in the mouse visual cortex enabled by tissue-level flexible mesh electronics with hexode arrays. Mesh electronics integrate seamlessly with neural networks, eliminating electrode drift. Action potentials from the same single units can be extracted through spike sorting and confirmed through waveform similarity and single-unit centroid position estimation evaluations. **b,** Schematics of chronic recording: electrical recordings were conducted on the mouse watching the same natural movie at different frequencies over months. The dynamics of neural representation were analyzed at both individual neuron and population levels over months. **c**, Bright-field (BF) microscopic image showing mesh electronics with five hexode arrays. **d**, Zoomed-in view showing individually addressable electrodes in each array. **e,** Representative photograph showing the mesh electronics implanted into the mouse brain with minimal acute tissue damage. The dashed line highlights the flexibility of the mesh electronics.

## Results

### Tracking natural movie stimulus-dependent representational drift from the same neurons over months

We fabricated the hexode flexible mesh electronics to record single-unit activities in head-fixed behaving mice using previously reported methods^15^ (Fig. 1c-d, Extended Data Fig. 1). Each mesh electronics consisted of five arrays of six highly packed electrodes with an interelectrode distance of 30 μm within each array (Fig. 1c-d). This distance is similar to the size of a single-neuron soma^16,17^, allowing each neuron to be simultaneously recorded by multiple nearby electrodes. We implanted the mesh electronics in the primary mouse visual cortices (Fig. 1e). The input/output (I/O) connectors of mesh electronics were connected to a lightweight flexible flat cable (FFC), enabling long-term stable electrical recording. The average interval of consecutive signal recording ranged from 3.4 ± 5.08 to 32 ± 21.3 (mean ± std) days from awake, head-fixed mice.

To consistently identify the electrical signals from the same neurons throughout multiple months of recording, we evaluated single-unit recordings based on their spike waveforms and estimated positions (Fig. 2a-b). For clarity, we refer to a unit as a putative neuron yielded by spike sorting throughout this discussion. We employed two criteria to consistently identify neurons over extended periods: high within-unit waveform similarity and minimal displacement of estimated unit centroid position across consecutive recording sessions. Specifically, we do not consider a neuron as consistently recorded across recording sessions if its within-unit waveform similarities were not greater than its similarities compared to other units (Fig. 2a). To estimate the centroid position of a single unit, we calculated a spatial average using electrode positions weighted by the square of the mean waveform amplitude collected at each neighboring electrode within a hexode^6^. This estimation is inferred by amplitude and is termed estimated single-unit centroid position. We then gauged the displacement of a unit by calculating the Euclidean distance between its estimated centroid positions across recording sessions. Notably, due to the complex morphology and biophysics of neurons, the estimated single-unit centroid position is unlikely to precisely match the physical location of neurons in relation to the electrode arrangement. However, if the signals remain stable and consistently recorded from the same neuron, the displacement of the estimated single-unit centroid position should also be negligible. In our study, units displaying greater than 10 μm of displacement (smaller than the size of the soma regions of one neuron) across two consecutive recording sessions were not considered consistent or stable (Fig. 2b).

**Fig. 2.**
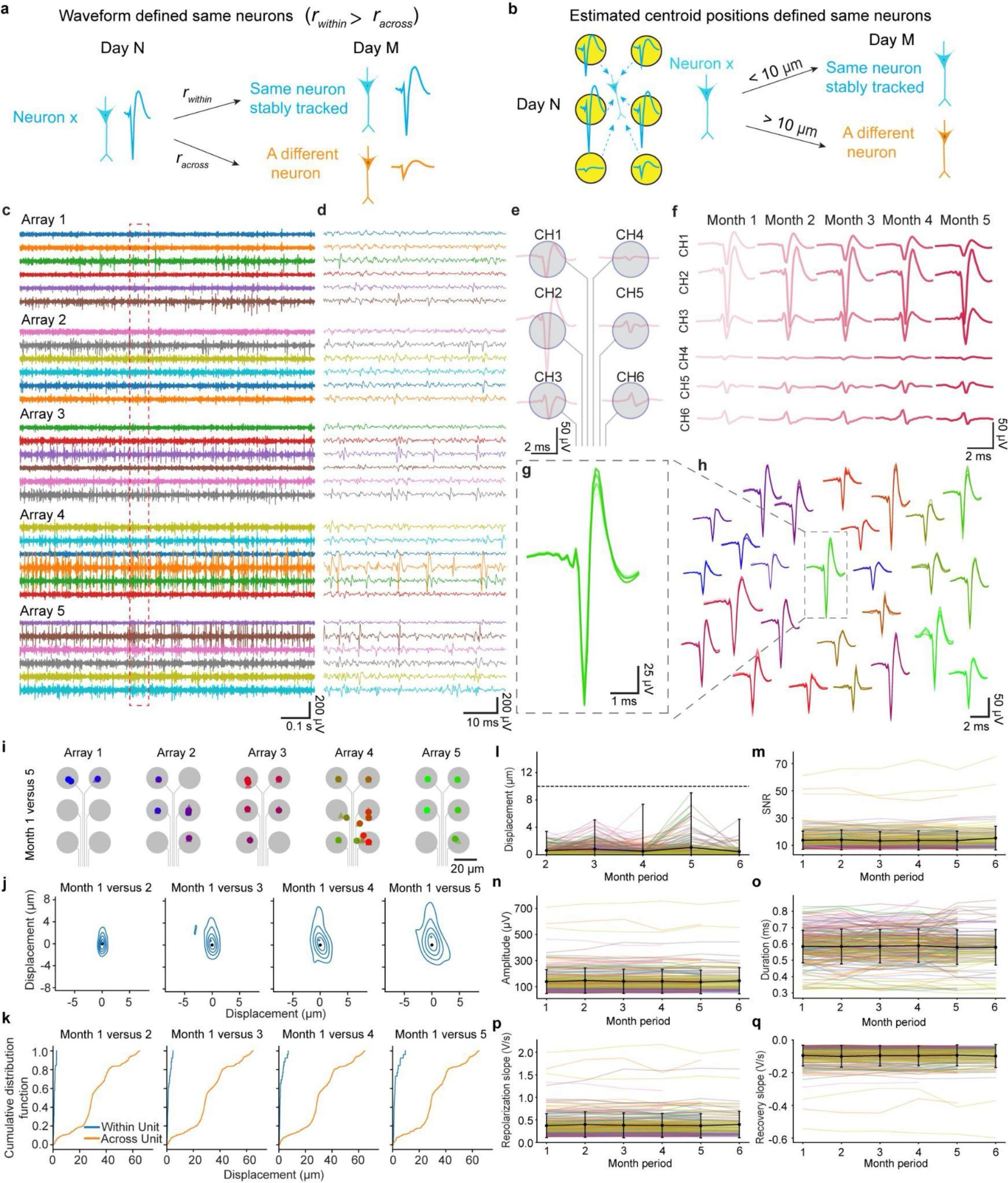
Stable tracking of neural activities from the same neurons over five months. **a-b,** Stable neurons defined by waveform similarity (**a**) and estimated single-unit centroid position displacement (**b**). **c,** Representative filtered voltage traces (Bandpass filter at 300 - 3000 Hz) showing spike dynamics recorded from representative hexode mesh electronics with five individual electrode arrays in an awake, head-fixed mouse at 1-month post-implantation. **d**, Zoomed-in view of the region (0.05 s) labeled by the red box in (**c**) showing action potentials simultaneously captured across different electrodes. **e**, Representative average waveforms recorded simultaneously at each of the recording electrodes of hexode mesh electronics. **f**, The waveforms from different electrodes recorded from each month plotted in order of gradient color. **g-h**, Superimposed waveforms from one representative neuron (**g**) and all neurons (**h**) on the electrodes with the highest amplitudes tracked over time. **i,** Estimated centroid position of single-neuron on month 1 and month 5 from all five electrode arrays in a representative mouse. The estimated centroid position for each single neuron on month 1 (triangle) was compared to month 5 (circle). Gray circles indicate the positions of the individual electrodes within an array. **j**, Mean displacement of estimated single-neuron centroid position from the mouse in **(i)** between month 1 (black dots, defined at origin) and months 2-5 (blue dots; columns 1–4, respectively). Contours indicate the quintile distribution of estimated single-neuron centroid position displacement for the neurons. **k**. Cumulative distribution of within-neuron estimated centroid position displacement (blue) between month 1 and months 2-5 (columns 1–4, respectively) and across-neuron centroid distance within each recording session (orange) for the mouse in (**i)**. **l,** Time-course analysis of the estimated single-neuron centroid position displacement between two consecutive months across all mice (*n* = 214 neurons from 9 independent mice). The dashed line indicates a displacement of 10 μm, a size smaller than neuron soma. **m,** Signal-to-noise ratio (SNR) of electrodes across recording sessions in each month. **n-q,** The mean waveform features – amplitude (**n**), duration (**o**), repolarization slope (**p**), and recovery slope (**q**) – of all neurons (*n* = 214 neurons from 9 independent mice) as a function of time. For (**l-q**), due to the difference in total length of the experiment in different mice, not all month periods have the same amount of data points.

To identify the signals from the same neurons and examine the stability of the signals, voltage traces were first filtered by 300–3,000 Hz bandpass filter and common referenced (Fig. 2c-d). We employed a fully automated approach (MountainSort4) to process the recording data concatenated from different recording sessions^18^. Subsequently, we conducted manual curation to extract the single-unit spikes across all recording sessions, employing the criteria described previously. Waveforms show that the spikes from the same neurons can be simultaneously detected by multiple nearby hexode electrodes, each showing varied amplitudes and spike shapes (Fig. 2e). The waveform shapes of the same neurons remained consistent throughout months of recording (Fig. 2f). Superimposing waveforms of individual neurons showed minimal variation during extended recording sessions (Fig. 2g-h). Single neuron waveforms were more similar with themselves across recording sessions than when compared to waveforms of any other neurons simultaneously recorded within a session/day. The estimated centroid positions of neurons remained nearly constant over several months, with displacements of identified neurons all under 10 µm across consecutive recording periods (Fig. 2i-l). Notably, the average distance between different neurons is greater than the displacement from the same neurons (Fig. 2k). Comprehensive statistical analyses across various mouse samples confirmed sustained signal-to-noise ratio (SNR) on different electrodes and similar waveform features in the same neurons recorded over time (Fig. 2m-q). The constant SNR demonstrated that the interface of the hexode electrodes to the recorded neuron did not degrade significantly throughout the entire recording period. Additional biological replicates showing stable neurons can be found in Extended Data Figure 2. In conclusion, our approach enables the isolation of the same neurons stably tracked across multiple months, with a total of 214 stable neurons over a maximal span of >5 months from 9 mice.

We presented mice implanted with flexible mesh electrode arrays in the primary visual cortex with 20 trials of a 30-second natural movie (three versions of the movie were used: 0%, 50%, and 100%, with 0% being the original movie and 100% being a noisy visual stimulus^8^; also see Methods) in a single stimulus session. Electrophysiological recordings were performed in a subset of the stimulus sessions, which we refer to as recording sessions, and were used for data analysis (Fig. 3a). The same neurons were consistently recorded based on the waveform shape and estimated neuron centroid position as described above (Fig. 2). We then computed the firing rate by binning the spikes for each stable neuron. Representative stable neurons with stimulus-evoked visual response can be found in Extended Data Fig. 3. We further qualitatively examined the movie stimulus-dependent representational drift of these stably recorded neurons by constructing a peristimulus time histogram (PSTH). Representative neural responses showed diverse responses to the natural movie stimulus over the entire course of the study (Fig. 3b-c). For instance, one representative neuron maintained a relatively consistent firing pattern throughout multiple movie repeats over different recording sessions spanning multiple months (Fig. 3b-c left). In contrast, another showed a directional shift of the firing pattern in response to movie repeats from different recording sessions (Fig. 3b-c middle). A separate representative neuron gradually developed a tuning pattern toward specific time bins in the natural movie over time (Fig. 3b-c right).

**Fig. 3.**
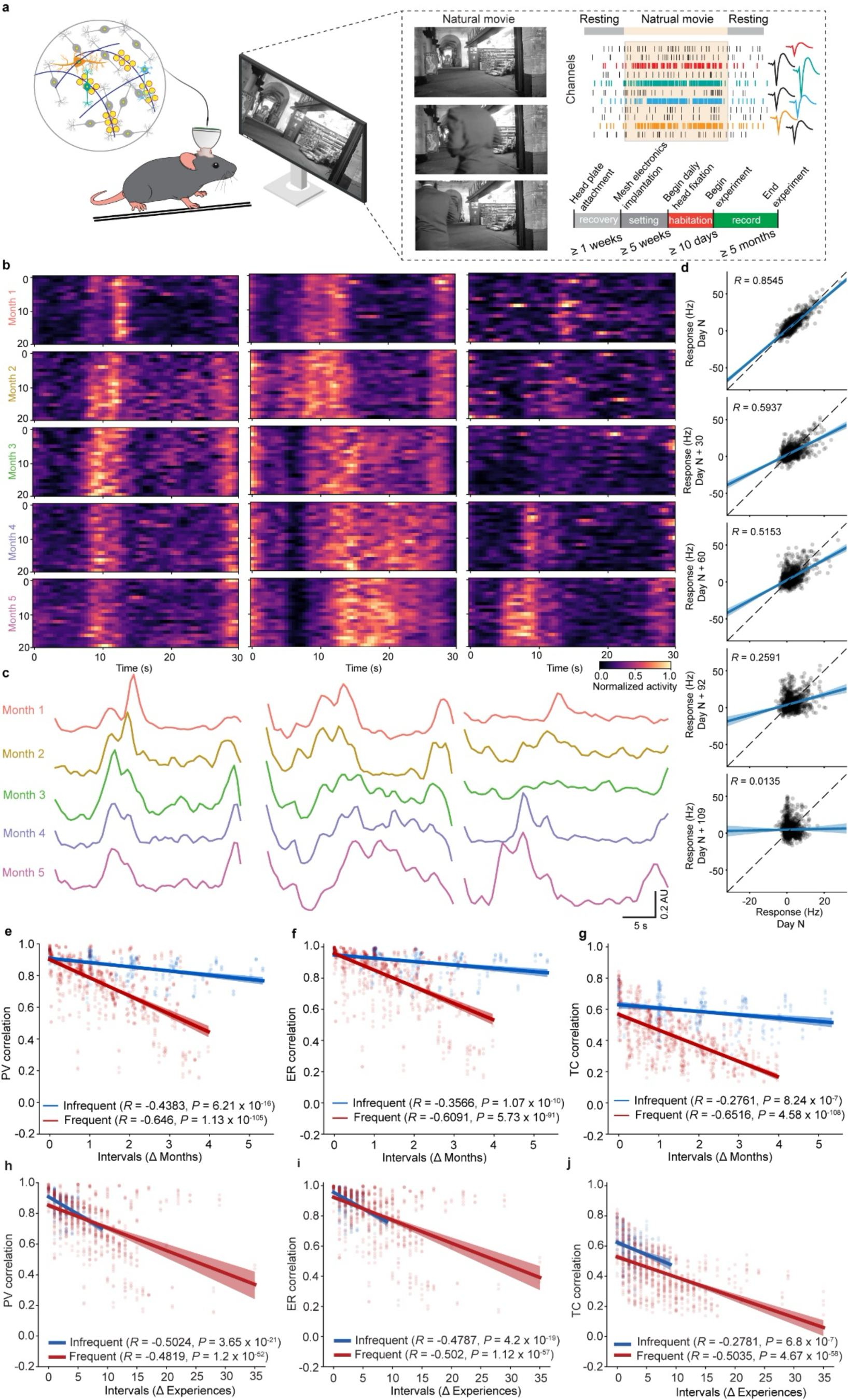
Representational drift at the level of the same individual neurons over multiple months. **a**, Experimental setup for the visual stimulation task. The visual cortex of the mouse was implanted with flexible hexode-like mesh electronics for long-term stable recording during the display of 30-second natural movies after animal habituation. Spikes from the same single units were sorted and curated for downstream analysis. **b**, Heatmap of the normalized firing rate from three representative neurons from one representative mouse stimulated frequently over the 5-month period. Each heatmap represents a neuron’s response to 20 repeats of the movie in a specific recording session, with each row in a heatmap representing an individual repeat of the movie. **c**, Mean of the normalized activity of the neurons in **b** across 20 repeats of the movie. Each line represents the neuron’s mean activity in a specific recording session. Scale bars correspond to a value of 0.2 AU of the normalized firing rate. **d**, Correlation between neurons’ firing rates at a former time (x-axis) and those at a later time (y-axis). Each dot represents one neuron’s mean firing rates at the two different times. The firing rate is baseline subtracted. The black dotted line is the unity line, and the blue line with shade is the regression line plotted with 95% confidence interval (CI). **e-g**, The correlations of PVs (**e**), ensemble rate vectors (**f**), and tuning curve vectors (**g**) between all pairs of recording sessions plotted against the corresponding time intervals for both groups of mice with frequent (*n* = 4 mice) and infrequent (*n* = 5 mice) visual stimulation. **h-j,** The correlations of PVs (**h**), ensemble rate vectors (**i**), and tuning curve vectors (**j**) between all pairs of recording sessions plotted against the corresponding experience number difference for both groups of mice. In (**e-j**), the straight lines with shades are the regression lines with 95% CI of the linear regression.

To quantitatively measure the representational drift in response to the same natural movie stimuli, we compared activities of each stable neuron in response to the same time bins across all pairs of recording sessions and found that the activities from the same neurons became less correlated across sessions with increasing time intervals (Fig. 3d). We then adopted metrics in existing literature^7^ to further quantify the dynamics of neural representation in V1 across sessions (Extended Data Fig. 3i): (1) the population vector (PV) representing the combined activity of a population of neurons and providing an overview of the collective activity patterns at individual time bins in the movie; (2) ensemble rate vector capturing the overall activity levels of the population of neurons during movie display; and (3) tuning curve vector capturing the changes in firing rate in single neurons with respect to the entire 30-second visual stimulus. By computing the correlations of population vectors (PVs) between different time bins in the movie across three consecutive days, we found high PV correlation between the same time bins across days, indicating the distinct neural coding of the movie sequence (Extended Data Fig. 3j). The same plot also exhibited similar correlation structures in recorded V1 neurons in the short term.

We computed the correlation of PVs, ensemble rate vectors, and tuning curve across all pairs of recording sessions (spanning >5 months). Decreasing correlations in each of the three vectors were observed across sessions as a function of increasing the time interval (Fig. 3e-g) in two groups of mice, one with more frequent (< 8 days/movie exposure) movie stimulation and the other with more infrequent stimulation (> 15 days/movie exposure). Extending previous short-term studies^7,8^ (< ∼2 months), this result suggests that representational drift in V1 neuron activities in response to the same movie stimuli can still be observed after monthslong period.

### Testing the hypothesis about the potential origin and mechanism of representational drift

We investigated several aspects of representational drifts in V1 neurons. Firstly, representational drift may be confounded by factors beyond the visual stimulus itself, such as pupil dynamics (an indication of animal arousal state) and signal instability^8,19,20^. To investigate this, we estimated the ratio of the pupil and eye at different times and found that change in the ratio remained consistent over time, suggesting no significant change in pupil dynamics (Extended Data Fig. 4a-c). Furthermore, we evaluated additional factors^6^ including waveform similarity, estimated neuron centroid position displacement, and changes in mean firing rates, as well as their relationship to tuning curve correlation across all recording sessions. Despite the varying levels of correlations in tuning curves, waveform similarity remained consistently high, displacement remained low, and changes in firing rate remained close to zero in both groups of mice (Extended Data Fig. 4d-i). These suggest that external factors associated with signal instability, probe instability, and pupil dynamics cannot fully explain the representational drift at the neuron level.

Secondly, previous reports have shown that neural activities in the piriform cortex stabilize upon more frequent stimulus, suggesting a stabilizing memory system under constant odor exposure^6^. It has also been reported that the visual cortex functions as a working memory system^21^. Therefore, we hypothesized that stimulus frequency may influence neural representation stability. Comparing two groups with different stimulus frequencies, we observed that, given the same time interval, neural activities in mice exposed to frequent stimuli drift more than in mice exposed to infrequent stimuli, as reflected by the greater drops in correlations of PVs, ensemble rate vectors, and tuning curve vectors in the frequent stimulus group (Fig. 3e-g). Notably, single neuron tuning exhibits lower correlation across various time intervals than PVs and ensemble rate vectors for both groups of mice (Fig. 3e-g). We then asked whether this difference was due to varying recording stability between the two groups. To test this, we examined various metrics for both groups. The results show that, in both groups, the within-neuron correlations were high (*r* > 0.9) (Extended Data Fig. 5a, c), and only small portions (< 10%) of the interspike intervals (ISI) were smaller than 1.5 ms (ISI violation^15^) (Extended Data Fig. 5b, d). ISI violation rates suggest that all neurons were expectedly constrained by the limits imposed by refractory period. Between the two groups, there was no statistically significant difference in within-neuron correlations, ISI violation rates, total number of stable neurons, or the percentages of stable neurons in the final recording sessions out of all units (Extended Data Fig. 5e-h). Changes in waveform features (duration, amplitude, peak-trough ratio, repolarization slope, and recovery slope) across the entire monthslong period were minimal and showed no significant difference between groups (Extended Data Fig. 5i-m). Taken together, recording quality cannot explain the differing drift amounts.

Thirdly, a recent study in hippocampus neural representation showed that elapsed time and elapsed experience number differentially drive the dynamics of neuron’s firing rate and spatial tuning, respectively^22^. We tested how drift changes with respect to the number of stimulus experiences received by mice. When studying the relationship between PV/ensemble rate vector correlation and the number of elapsed stimulus sessions, we identified similar correlation change for both groups of mice (Fig. 3h-i, Extended Data Fig. 6a-b). The change in the correlation of single neuron tuning was less consistent between the two groups (Fig. 3j, Extended Data Fig. 6c), suggesting that elapsed experience could not fully explain the extent of drift in single neuron tuning activities in V1. Overall, changes of V1 neural representation are largely influenced by the number of experiences rather than time elapsed alone.

Fourthly, we examined temporal dynamics of drift to understand how stimuli influence the dynamics of neural representation over time. By calculating the cosine distance between PVs at the same movie time points across consecutive recording sessions, we observed a non-decreasing trend in both drift amount and rate for both groups of mice (Extended Data Fig. 6d-e), despite a higher drift rate in PVs in mice exposed to frequent stimulation. We divided the recording sessions in the monthslong period into two halves to investigate PV drift rate. Specifically, we computed the drift rate across all pairs of recording sessions in each half. We found that, in both groups of mice, drift rates in the first half were not significantly different from those in the second half (Extended Data Fig. 6f). This confirms no significant decrease in the drift rate of PVs in V1 over time for both groups of mice, thereby indicating that no stabilizing neural representation can be inferred.

Fifthly, given substantial drift in single-neuron tuning in response to the movie stimulus (Fig. 3g, j), we asked whether the response geometry is conserved (Extended Data Fig. 7a). A previous study suggested that drift could be attributed to population geometry rotation^6^. To test this hypothesis, we investigated visual response geometry dynamics by measuring angles formed between two edges connected by a single vertex (see Methods) and discovered that the visual geometry remained relatively stable in mice with infrequent stimuli but changed more in those mice with frequent stimuli (Extended Data Fig. 7b-d).

Finally, we asked whether higher-order statistics in visual stimulus affect representational drift in neurons. In a previous study, movies were parametrically phase-scrambled to three levels: 0%, 50%, and 100%, and visual responses were recorded for two months using calcium imaging^8^. The results showed that the extent of parametric scrambling did not produce a significant trend in representational drift. We examined whether long-term electrophysiological data from the stable neurons would yield similar results. We present 20 repeats of the 50% and 100% phase scrambled movie variants to the mice (Extended Data Fig. 8a) and then quantified the stability of single-neuron tuning towards the same movies across different recording sessions. In both groups of mice, neuron tuning in response to the original movie was statistically more correlated over time than turning in response to 50% and 100% phase scrambled movies (Extended Data Fig. 8b-d, f). When calculating the drift rate of the tuning curve vectors, we observed that those in response to the original movie were significantly higher than those in response to 50% and 100% variants of movies for both groups of mice (Extended Data Fig. 8e, g). These findings suggest that representational drift is dependent on stimulus type and their higher-order statistics with possible distinct short- and long-term dynamics.

### Tracking latent dynamics changes on the same neural manifold over months

In both groups of stimulus frequency, single neurons’ tuning to the movie became unstable (Fig. 3g, j). The visual response geometry to the movie was not maintained, especially in mice exposed to frequent stimuli (Extended Data Fig. 7). Previous studies showed that neural correlates of the external world, such as motion intent, can be embedded in a population-level low-dimensional latent space termed neural manifold^23^. We therefore ask whether a long-term stable structure on the neural manifold can emerge from these drifting neural representations.

To uncover time-varying neural latent dynamics in a low-dimensional neural manifold, we combined neural activities from all recording sessions and applied a Gaussian-process factor analysis^24^ (GPFA) (Fig. 4a). GPFA is a linear dimension reduction method specifically designed for uncovering the low-dimensional latent dynamics from trial-based neural activity data. We performed GPFA on the data from all mice. Representative results from one mouse in each frequency group were shown in Fig. 4b-e. In the mice with frequent stimulation, plotting the top two dimensions revealed that the smoothed latent dynamics averaged across all repeats in different recording sessions showed significant changes (Fig. 4b, d). For these mice, the correlation of latent dynamics (top three dimensions) between sessions declined as time intervals between sessions increased (Fig 4f top, g, h left). This decline was less evident in mice receiving less frequent stimuli (Fig. 4f top, g, h right). Notably, these correlation changes in latent dynamics were better explained by elapsed experience numbers (Fig. 4f bottom).

**Fig. 4.**
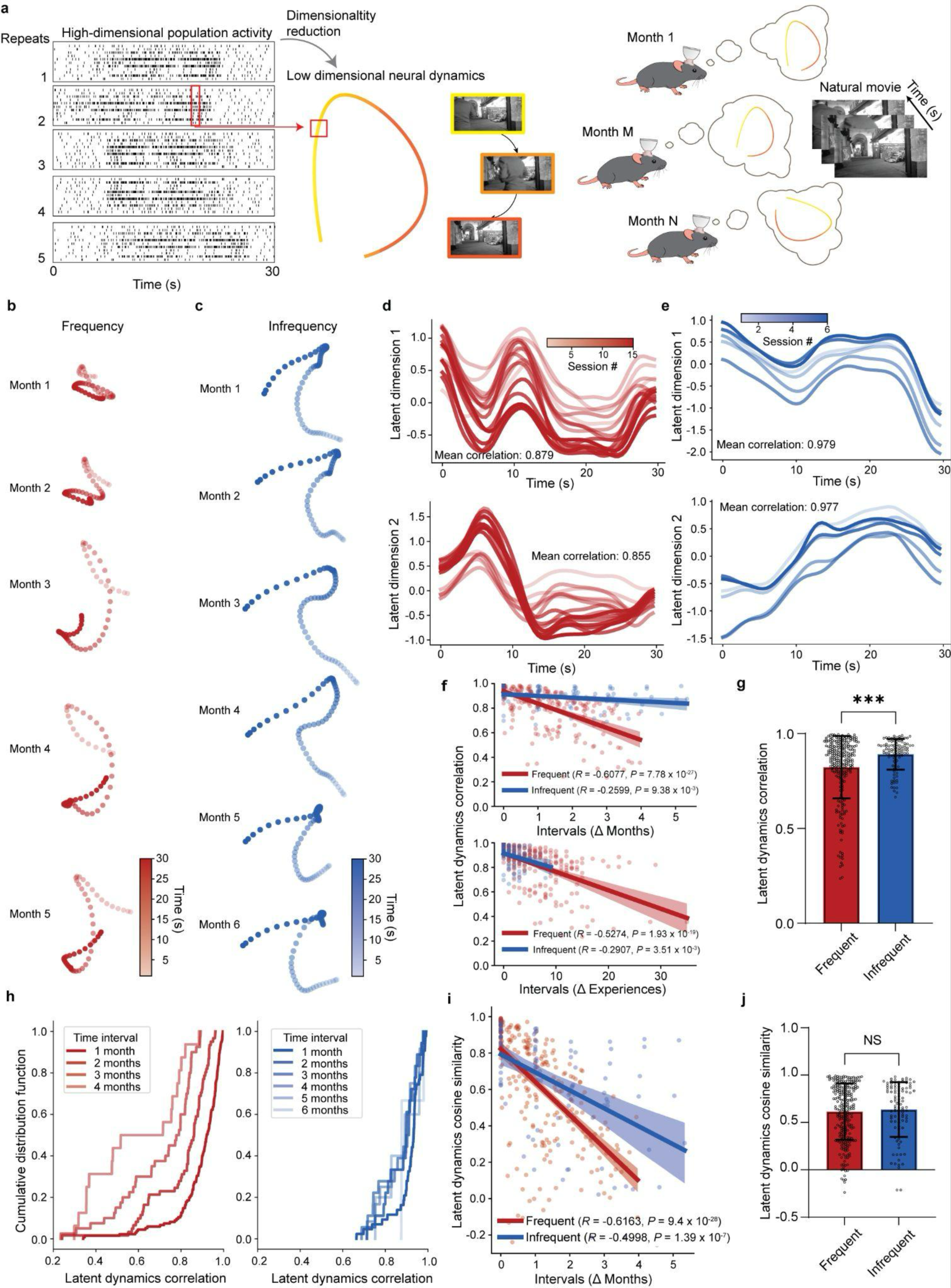
Neuron population latent dynamics in response to the same visual stimuli over multiple months. **a,** Extraction of latent dynamics (low-dimensional neural trajectory) from neural activities in response to the visual stimuli in different recording sessions across months using Gaussian process factor analysis (GPFA). One single GPFA was fitted to the data combined from all recording sessions in each mouse in this figure. **b-c,** The smoothed latent dynamics averaged across all movie repeats in each recording session over multiple months from one representative mouse with frequent visual stimulation (**b**) and another with infrequent visual stimulation (**c**). **d-e,** The latent dynamics projected onto the first (top) and second (bottom) latent dimensions for representative mice with frequent (**d**) and infrequent (**e**) visual stimulations. Values indicate the mean correlations between all pairs of the latent dynamics on a specific latent dimension. **f**, Top: averaged correlations of latent dynamics across the top three latent dimensions between pairs of sessions plotted against the time interval, for all mice stimulated frequently and infrequently. Bottom: same as top but plotted against the difference in experience number. **g,** Bar plot showing the mean, standard deviation, and distribution of the pooled averaged correlations of latent dynamics between all pairs of sessions for both mouse groups (*P* = 0.0008; two-tailed Mann-Whitney test; frequent: *n* = 252; infrequent: *n* = 99). **h,** CDF of averaged correlations of latent dynamics, grouped by different ranges of intervals, for both mice stimulated frequently (left) and infrequently (right). **i,** Averaged cosine similarity of latent dynamics across the top three latent dimensions between pairs of sessions plotted against the time interval, for all mice stimulated frequently and infrequently. **j,** Bar plot showing the mean, standard deviation, and distribution of the pooled averaged cosine similarity of latent dynamics between all pairs of sessions for both mouse groups (*P* = 0.618; two-tailed Mann-Whitney test; frequent: *n* = 252; infrequent: *n* = 99). In **f** and **i**, the regression lines are plotted with 95% CI.

Further examination of the mean latent dynamics across different recording sessions revealed a translation or shift on the same neural manifold (Fig. 4d-e). We then used cosine similarity to compare latent dynamics across different recording sessions. Mathematically, unlike Pearson’s correction, cosine similarity does not center variables, making it sensitive to shifts and translations (see Methods). In both groups, cosine similarity between the latent dynamics rapidly declined with increasing time intervals between recording sessions (Fig. 4i). The difference between Pearson’s correlation and cosine similarity (Fig. 4f-g, i-j) highlights the translation of the latent dynamics within the same neural manifold over time.

Finally, we explored whether latent dynamics in response to the three variants of the movies on the same neural manifold exhibits significant difference in cross-session correlation (Extended Data Fig. 8a). Applying GPFA to the remaining two variants of the movie, we compared latent dynamics between all session pairs. Results showed that latent dynamics under the original movie were statistically more correlated than those under parametrically scrambled movie stimuli (Extended Data Fig. 8h-k). Therefore, the stability of neural activity is affected by higher-order statistics both at the individual neuron and population levels.

### Transformation leads to more stable latent dynamics over months

The translation of the latent dynamics within the same neural manifolds across different recording sessions was suggested by the observation that Pearson’s correlation and cosine similarity yield divergent results (Fig. 4f top, i). To address this, GPFA was applied to neural activity data from each recording session, during which the neural activity data were translated through data centering. The resulting latent dynamics from different recording sessions resided on distinct neural manifolds. To compare the latent dynamics, these neural manifolds were aligned by linearly transforming the loading matrices obtained by GPFA across consecutive recording sessions^11^. This is referred to as the translated stage. After alignment, the vectors defining the top three dimensions, termed neural modes, were highly similar across different times for both groups of mice (Fig. 5a; Extended Data Fig. 9a-d). No significant difference in alignment results was observed between mice stimulated frequently and those stimulated infrequently (Fig. 5b). Notably, in more frequently stimulated mice with less correlated latent dynamics, we were able to identify latent dynamics that are more similar when using cosine similarity as the metric, attributed to the translation and centering of the data (Fig. 5c-f). Note that onwards we mean high cosine similarity as “similar” unless specified otherwise. Compared to those before translation, correlations of the latent dynamics after translation across pairs of recording sessions were compromised in mice stimulated frequently (Fig. 5g-h).

**Fig. 5.**
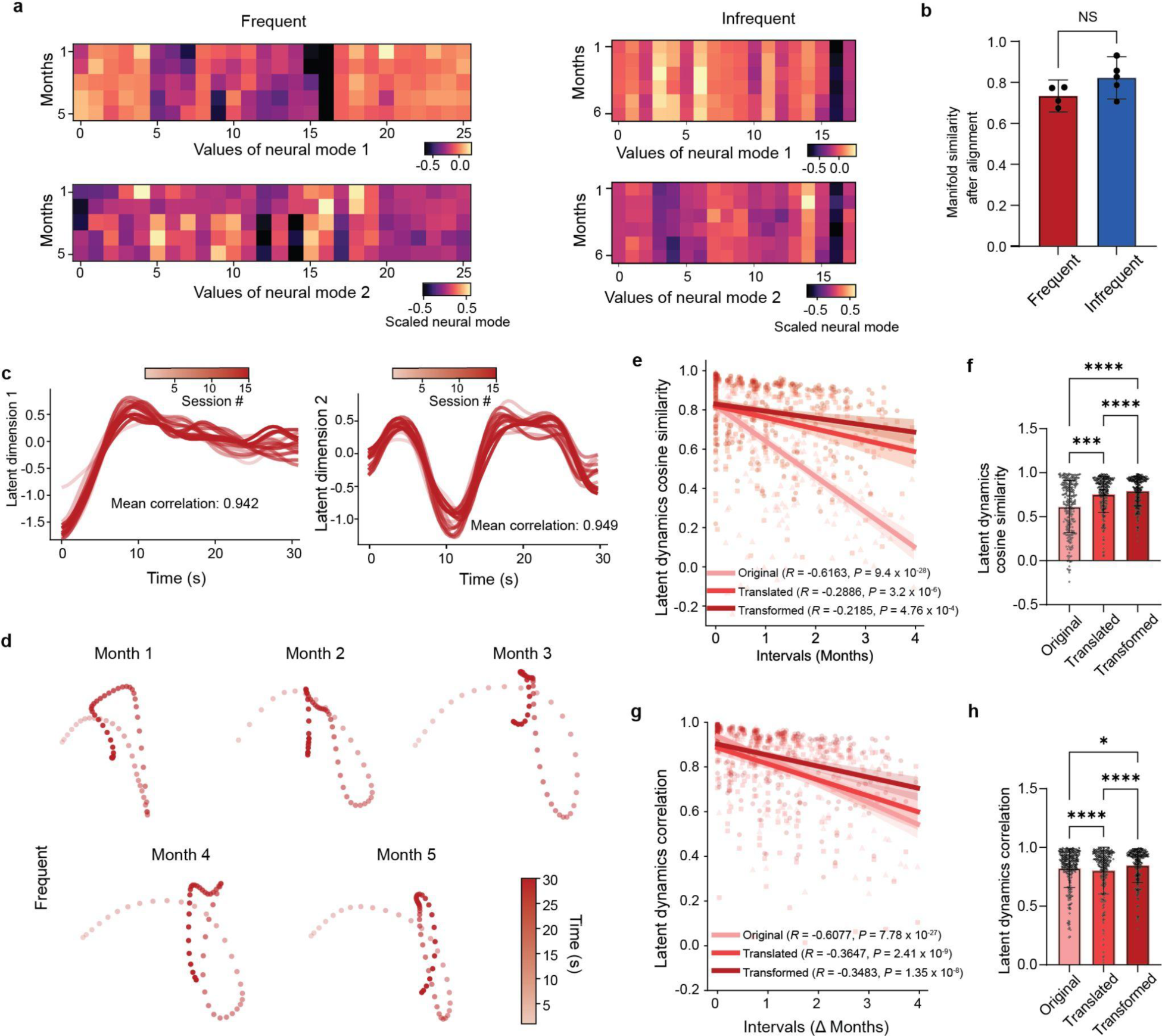
Highly correlated and similar latent dynamics over multiple months after translation and transformation. **a,** The neural modes (vectors defining each axis of the neural manifold) in different recording sessions for the first (top) and second (bottom) latent dimensions after translation of latent dynamics and alignment of neural manifold, for both mice stimulated frequently (left) and infrequently (right). Each row in the blocks is a vector representing a neural mode. **b,** The averaged cosine similarity of all five neural modes across different pairs of recording sessions for both groups of mice (*P* = 0.1056; two-tailed unpaired t-test; frequent: *n* = 4; infrequent: *n* = 5). **c,** The latent dynamics after transformation projected onto the first (left) and second (right) latent dimensions for representative mice with frequent visual stimulations. Values indicate the mean correlations between all pairs of the latent dynamics on each latent dimension. **d,** The averaged smoothed latent dynamics across all movie repeats after transformation in each recording session over multiple months for one representative mouse with frequent visual stimulation. **e**, The averaged cosine similarity of latent dynamics between pairs of recording sessions before (original, same as in Fig. 4) and after latent dynamics translation and manifold alignment (translated), and after transformation (transformed), plotted against the different time intervals for mice stimulated frequently. **f,** Bar plot showing the mean, standard deviation, and distribution of the cosine similarities of latent dynamics between all pairs of recording sessions at the three stages of transformation (original vs translated: *P* = 0.0001; original vs transformed and translated vs transformed: *P* < 0.0001; Friedman test and Dunn’s multiple comparison test). **g**, Same as in **e** but for the averaged correlations. **h**, Same as in **f** but for the averaged correlations (original vs translated and translated vs transformed: *P* < 0.0001; original vs transformed: *P* = 0.0106; Friedman test and Dunn’s multiple comparison test).

The reduced correlation of latent dynamics suggests a change in the shape of the latent dynamics across different times. We then asked whether a linear transformation of the latent dynamics across recording sessions could estimate more similar and correlated latent dynamics for encoding the same visual stimulus in the long term. After transforming the latent dynamics, we found that, in the mice with more frequent stimuli, both the correlation and the cosine similarity of the latent dynamics were significantly greater than those characterized before translation and transformation (Fig. 5e-h). The effect of translation and linear transformation on latent dynamics for representative mice with infrequent stimulation and additional mice with frequent stimulation is shown in Extended Data Fig. 9. Although translation substantially increased the cosine similarity of the latent dynamics over time, especially in frequent mice, it did not capture potential transformation of the latent dynamics undertaken by the neuron population. To identify a more similar and correlated structure, the time-varying latent dynamics were linearly transformed, constituting an affine transformation.

Ultra-flexible hexode electrode arrays can provide a stable interface between the electrodes and recorded neurons. However, we questioned whether it is still possible to align the drift with an affine transformation when this drift includes both representational drift and recording instability such as probe drifting, signal degradation, and eventual drop out (Extended Data Fig. 10a). To assess this, we stimulated recording instability within the neural data (see Methods). After conducting multiple simulation at different percentages of probe instability and applying the same analytical steps as before (GPFA, translation, and transformation of the latent dynamics), we computed mean correlations of the latent dynamics. This analysis utilized two example mice from each stimulus frequency group, with results showing that as the percentage of probe instability increased, the mean correlations of transformed latent dynamics decreased (Extended Data Fig. 10b-g). Consequently, when intrinsic neural representational drift is compounded with recording instability, it becomes more challenging to identify a stable neural correlate over time that corresponds to the same visual information. This further suggests the importance of using long-term stable BMIs, such as the flexible electrodes, in research on neural representations and in developing stable neural decoders.

### High performance decoding from a single visual information decoder over months

Given the identification of consistent low-dimensional latent dynamics through affine transformation (Fig. 6a-b), we asked whether a single machine learning-based decoder, trained for each mouse, could reliably and continuously decode visual information stably recorded from the same neurons in brain regions with intrinsic representational drift. As an example, we trained a decoder that can take in windows of neural data as input and predict the time window to which the input data corresponds (Fig. 6b).

**Fig. 6.**
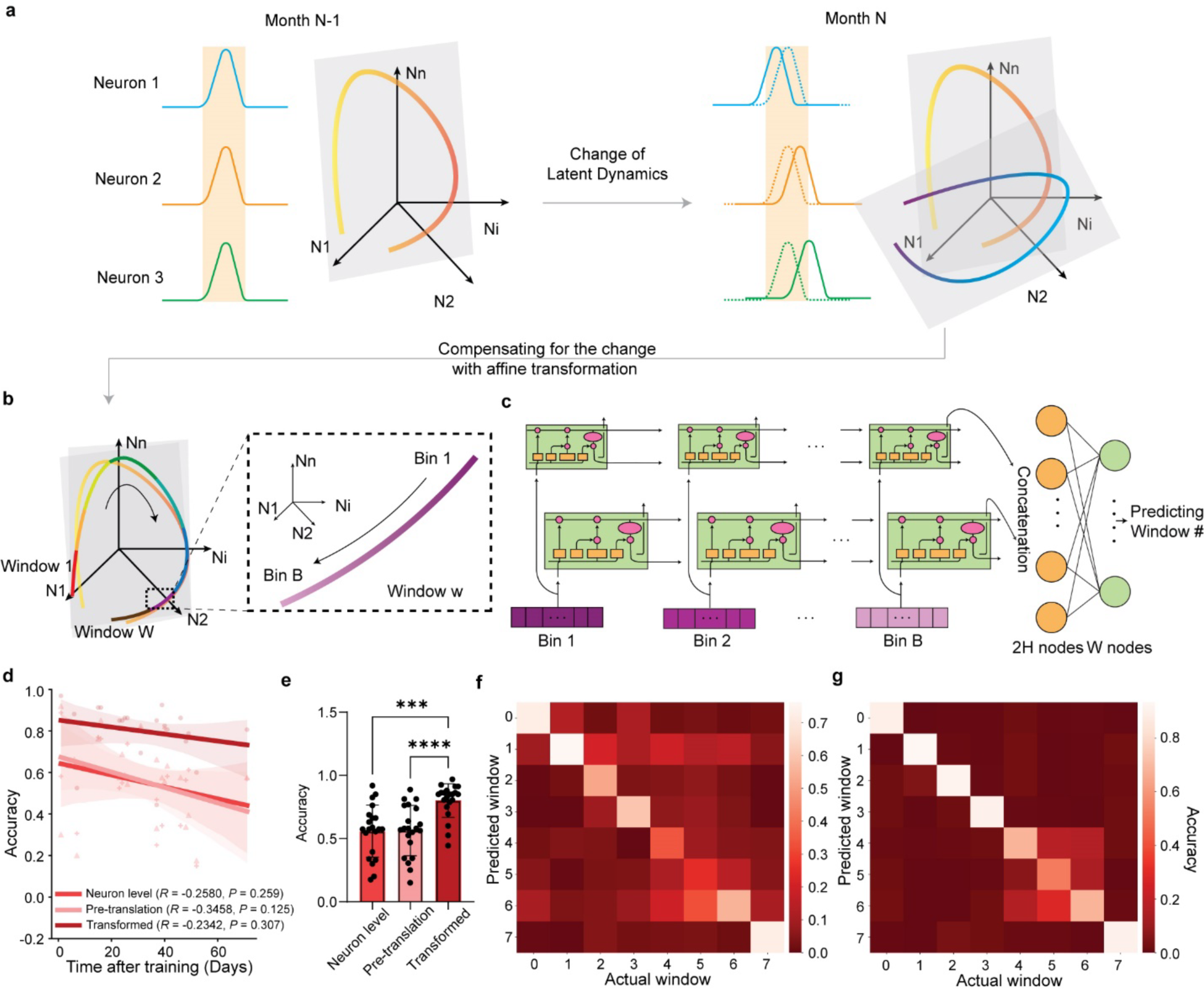
Reliable decoding of visual information from neural activities in a drifting brain region after transformation of neural population latent dynamics. **a,** Schematics illustrating the changes in single-neuron activities across different sessions towards the same visual stimulus, resulting in changes in low-dimensional neural population dynamics and neural manifolds. **b-c,** Inputting the low-dimensional latent dynamics after transformation (**b**) into a neural network with a long short-term memory (LSTM) architecture and a densely connected linear layer (**c**). The LSTM sequentially takes in latent dynamics from one time bin at a time to predict the time window in the movie. There are W time windows (W = 8) for the 30-second movie. Each time window has a sequence of B time bins (B = 15; 0.25-second for each bin). Two layers of LSTM produce two output vectors (each of dimension H) at the output gates, and concatenation of the two outputs is passed into a densely connected linear layer with W output nodes that are used to predict the corresponding time window. **d,** The accuracy of visual information decoding from neural activities in different sessions with one single trained LSTM decoder for mice with frequent visual stimulation. The inputting data are high-dimensional single-neuron activities (neuron level), latent dynamics before translation (pre-translation), and latent dynamics after transformation (transformed). **e,** The comparison of decoding accuracies at different stages in **d** (transformed vs neuron level: *P* = 0.0002, transformed vs pre-translation: *P* < 0.0001; Friedman test and Dunn’s multiple comparison test; n = 21 pairs). **f-g,** Confusion matrices of time window predictions from neuron-level activities (**f**) and latent dynamics after transformation (**g**), for mice with frequent stimulations.

Firstly, we built a decoder based on a recurrent neural network (RNN) with a long short-term memory (LSTM) architecture. This decoder was designed to sequentially process neural data at individual time points within a given time window and predict the identity of that window. The architecture consisted of two LSTM layers followed by a densely connected linear layer (Fig. 6c). We split all recording sessions into two halves based on the time of recording within the monthslong recording period: an earlier half and a latter half. The windows of neural data from the earlier half were used to train the decoder in a supervised manner, while the decoder’s performance was evaluated by predicting the windows of neural data from the latter half. The decoder used three types of input data: individual neuron firing rates, latent dynamics before affine transformation, and latent dynamics after affine transformation. Training and evaluating the decoder with each of these three input types showed that for mice with frequent visual stimuli, using the transformed latent dynamics as input data significantly improved prediction accuracy compared to using individual neuron firing rates or untransformed latent dynamics as inputs (Fig. 6d-g).

To validate these results, we used another decoder, a support vector machine (SVM) classifier, to train and evaluate the neural data. Unlike the sequential input method used by the LSTM, the SVM used concatenated neural data from all time points in one time window as input. We then evaluated the prediction accuracy over multiple months using the three types of input data. The results obtained with SVM were consistent with those from the LSTM (Extended Data Fig. 10h-k). Performance of the two decoders for the mouse group with infrequent stimulation are also shown in Extended Data Figure 10. Taken together, transformed latent dynamics in response to the same visual stimuli were preserved over multiple months, allowing a single decoder to consistently predict temporal information in visual stimuli encoded in intrinsically drifting neural activities with a high performance over multiple months.

## Discussion

In this study, we integrated tissue-like, flexible hexode electrode arrays with a specifically designed software pipeline for the consistent selection and tracking of the same neurons over months. Using waveforms from multiple nearby electrodes and estimated neuron centroid positions, we tracked visual stimulus-dependent, single-neuron action potentials from 214 stable neurons across 9 mice for up to >5 months. In line with previous studies^7,8^, we identified neural representational drift in the mouse visual cortex when exposed to natural movie stimuli. However, unlike previous studies that focused on the representational drift over days or weeks, the use of flexible electronics for neural recording allows for precise characterization of this phenomenon over a monthslong period. This extended capability to characterize representational drift in the visual cortex has facilitated testing hypothesis about the origins and potential mechanisms of representational drift, observing latent dynamics transformation during neural representational drift, and developing strategies to overcome representational drift for stable brain information decoding by leveraging flexible brain-computer interfaces.

First, we tested hypotheses regarding the origins and mechanisms of representational drift in the visual cortex. Previous studies suggested that, in certain brain regions such as hippocampus, representational drift is associated with memory formation and consolidation, potentially as a byproduct of the continuous flux of new memories^6,20,25^. Moreover, the piriform cortex, known for its rapid learning and memory functions, demonstrates a history-dependent stabilizing representation of odor responses after repeated stimulus exposure, suggesting memory stabilization. The primary visual cortex is also suggested to function as a working memory system^21^. To test this, we divided mice into two groups based on stimulus frequency to investigate the potential role of the memory system as a driver of drift in the visual cortex. Our findings indicate that higher stimulus frequencies caused greater drift at the individual neuron level without stabilizing memory representations. In addition, our analysis revealed consistent drift rates throughout the experiment, suggesting a continuously evolving neural representation. We also found similar drift at the individual neuron level across the two groups of mice, related more to the number of experiences rather than the time elapsed, highlighting a dynamic representation system activated by each stimulus. While our data do not support the involvement of long-term memory formation in the visual cortex, they do not exclude the presence of a working memory system operating on a shorter timescale than our experiment. In addition, the widely observed representational drift in different brain areas is likely regulated by different mechanisms.

Next, we observed the instability of latent dynamics within neural population activity. Previous results identified a stable low-dimensional structure for approximately one month in the visual cortex using nonlinear dimension reduction methods such as tSNE and ISOMAP^7,26^. However, the dynamics of such a structure over longer terms remained elusive. In this study, we used a linear method, GPFA, to study representational drift in the visual cortex. This method yields interpretable latent dynamics and neural subspace, whose axes correspond to individual neuron weights. By applying GPFA to combined neural data from all recording sessions, we identified correlated latent dynamics (R ≈ 0.8) over a timescale comparable to a previous study^26^ (∼1.5 months). However, over longer periods, our results indicated a significant translation of the latent dynamics for both groups of mice with different stimulus frequencies. Notably, for mice with infrequent stimulation, although ensemble rate vectors from single neurons remained relatively constant over time, the linear combinations of neurons induced a shift in latent dynamics on the low-dimensional manifold.

Then, we applied GPFA individually to each session’s data, which involved data centering. Our long-term results in mice with frequent stimulation showed the latent dynamics gradually changed in shape. The latent dynamics from later sessions were therefore transformed based on earlier sessions. We argue that transforming the latent dynamics is equivalent to changing the neural manifold and thereby the weights of neurons. This transformation resulted in highly similar and correlated latent dynamics over the long term. This is particularly significant in mice exposed to frequent stimulation, suggesting that visual information can be stably encoded at neural population dynamics level in the visual cortex. For mice with infrequent stimulation, the population-level latent dynamics corresponding to the movie exhibited shift, but their shapes were highly correlated even before transformation. Translation makes the latent dynamics more similar. Kept minimally stimulated by the specific natural movie, neurons in the infrequent stimulation group maintained high stability in various levels such as latent dynamics shape, PVs, ensemble rate vectors, and visual response geometry, although it is important to note the less correlated single-neuron tuning and the shift of latent dynamics. Altogether, GPFA reveals drift as a model estimated by affine transformation, and after compensating for such changes, we can identify a more stable representation in response to the same visual stimulus across all mice for up to more than five months.

Our results also suggest that the neural network in the visual cortex is a dynamic system with constantly changing neural activity, unlike the gradually stabilizing representation observed in the olfactory cortex. Another study using Ca^2+^ imaging and Neuropixels characterized individual neuron-level drift and revealed less stable (less correlated) representations than our results on a similar timescale in V1^7^. Additionally, it was previously concluded that changing higher-order statistics in visual stimulus did not generate a significant difference in drift over a timescale of around one month^8^. However, we observed a notable trend in representation stability in response to differentially scrambled movies. These differences might be attributed to the possibility that different mechanisms drive drift across different timescales and brain regions. Variations in the sensitivity, resolution, and number of neurons analyzed using flexible probe^2^, rigid electrode arrays^9,27,28^, and optical technologies^29,30^ could also have influenced these observed differences. By simulating recording instability, we identified a significantly compromised ability to apply transformation to uncover stable and correlated latent dynamics. This suggests that probe instability is likely to confound research in neural representation, leading to largely divergent conclusions across studies. The inability to stably track activity from the same neurons makes it challenging to distinguish recording instability from the brain intrinsic dynamics. Therefore, further studies on drift phenomena may benefit from developing large-scale, 3D stacked, chronically stable neural interfacing technologies^31,32^ for potential stable tracking of a larger neuronal population under the same platform.

While our study proposed a linear model to estimate representational drift in the local neural network of V1, the underlying mechanisms and potential functions of representational drift across different brain regions are yet to be fully understood. Some theories suggest that representational drift could be an incidental outcome of the brain’s use of redundancy in its high-dimensional population coding to facilitate physiological functions^25^. Inspired by artificial neural networks, representational drift could also be part of computational features that regularize the system, such as dropout and noisy synaptic updates for optimization of functions related to memory and continual learning^33^. Future longitudinal studies utilizing large-scale flexible electronics to track more stable neurons could help resolve ongoing debates, such as the nature of drift in the hippocampus^5,34,35^ and the existence of drift in motor-related brain region^10,12–14,36–38^, thereby deepening our understanding of learning, memory processes, and neural computation^39–41^. Our technology can also benefit the long-term stable and accurate brain decoding in future applications. Traditional BMI applications have suffered from probe instability, significant neuron loss, as well as intrinsic drift in neuron activities^10,11,42^, requiring constant recalibration and stabilization. Here, the capability to track activities from the same neurons via flexible brain-computer interfaces could potentially lead to a neural decoder that more accurately characterizes representational drift in each brain region. This, in turn, could be used to estimate the drift and counteract its effects through processes such as mathematical transformation. Additionally, with improved neural signal stability, a broader range of information could be accurately decoded, significantly enhancing BMI performance. Our study demonstrates a long-term stable single BMI decoder that leverages similar latent dynamics based on a known stimulus sequence. However, a broader challenge remains: determining whether transformations can effectively capture drift in neural activities in response to various sensory stimuli, which could offer a universal framework for characterizing drift and brain decoding in V1 and other brain regions.

## Methods

### Device fabrication

Fabrication of the ultra-flexible mesh electronics with hexode array was based on methods described previously^15^. Key steps are described as follows. The 3-inch Si/SiO_2_ wafers were utilized as insulating substrates for the fabrication process of mesh electronics. The Si/SiO_2_ wafers were cleaned with acetone, IPA, and deionized (DI) water and were dried using nitrogen gas (N_2_). In order to enhance the adhesion of photoresists with the substrate, a spin-coating method was employed to apply hexamethyldisilazane (HMDS, MicroChem) at a speed of 4000 rotations per minute (rpm). Two photoresists, LOR 3A and S1805 (MicroChem), were subsequently spin-coated onto the Si/SiO_2_ wafers at the speed of 4000 rpm and baked at 180°C for 5 minutes and at 115°C for 1 minute, respectively. To expose the Ni sacrificial layer, a Karl Suss MA6 mask aligner equipped with 365 nm ultraviolet (UV) light was used to apply a dosage of 40mJ/cm^2^. The exposed layer was developed using CD-26 developer (MICROPOSIT) for 70 seconds. To remove any remaining photoresist residues, an O_2_ plasma (Anatech Barrel Plasma System) treatment was conducted with 50 watts of power for 30 seconds. A Sharon Thermal Evaporator was employed to deposit a 100-nanometer-thick layer of Ni. Afterwards, a standard lift-off procedure was conducted in remover PG (MicroChem) for 2 hours. Once the Ni layer was patterned, a precursor SU-8 (specifically SU-8 2000.5 from MicroChem) was spin-coated onto the wafers at 4000 rpm. The coated wafers were pre-baked at temperatures of 65°C and 95°C for 2 minutes each, followed by exposure to 365 nm UV light at a dosage of 200 mJ/cm^2^. After another round of baking (post-exposure baking) at 65°C and 95°C for 2 minutes each, the wafers were developed using SU-8 developer for 60 seconds. They were then rinsed with isopropyl alcohol (IPA) for 30 seconds, dried with an N_2_ gun, and hard-baked at 180°C for 40 minutes to define mesh-like SU-8 patterns with a thickness of 500 nm, serving as the bottom encapsulation layer. Following the patterning of the SU-8 bottom layer, the HMDS/LOR3A/S1805 photoresist layers were spin-coated using the same procedure as previously described. Subsequently, a 5/80/5-nanometer-thick layer of chromium/gold/chromium (Cr/Au/Cr) was deposited using an electron-beam evaporator (Denton). The standard lift-off procedure in remover PG (MicroChem) was then conducted overnight to define the Cr/Au/Cr interconnects. The same photolithography process was repeated to define 5/50-nanometer-thick chromium/platinum (Cr/Pt) layers as the electrodes. After the electrode patterning, the top SU-8 encapsulation layer was patterned using the same method as used for the bottom SU-8 layer. For the metal shuttle-based implantation, we designed a polymer hole on the top of the mesh electronics. For the polymer shuttle-based implantation, the SU-8 precursor (SU-8 2025, MicroChem) was spin-coated, pre-baked, exposed to UV light, and post-baked to form the SU-8 anchor patterns that connected the mesh and SU-8 shuttle. The wafer was spin-coated with dextran solution, and a subsequent layer of SU-8 precursor was applied with pre-heating and post-baking steps. Another layer of SU-8 precursor was spin-coated, following the same pre-baking and UV exposure steps, to define the SU-8 shuttle pattern. The developed patterns were then hard-baked to finalize the anchor and shuttle patterns.

### Animals and ethical compliance

Adult male C57BL/6 mice (Charles River Laboratories) at 16 weeks of age were used in this study. The mice were housed in an animal facility at 22 ± 1 °C with humidity ranging from 30% to 70% under a 12-hour light/dark cycle. The mice were fed ad libitum. All animal procedures complied with the National Institutes of Health guidelines for the care and use of laboratory animals. They were also approved by the Harvard University Institutional Animal Care and Use Committee under protocols 19-03-348 and 20-05-368.

### Surgical implantation

Device implantation followed previously established methods^15^. Briefly, to ensure sterility, all metal tools that came into direct contact with the animal subjects underwent sterilization with autoclave and bead-sterilization for 1 hour before use. Similarly, all plastic tools that had direct contact with the animal subjects were sterilized with 70% ethanol, rinsed with sterile DI water and sterile 1× PBS prior to implantation. Mice were anesthetized with 2–3% isoflurane through inhalation and maintained under anesthesia with 0.75–1% isoflurane. The level of anesthesia was verified through a toe pinch test before the commencement of surgery. The anesthetized mouse was positioned in a stereotaxic frame equipped with two ear bars and one nose cone. With a sterile scalpel, a 5 mm longitudinal incision was cut in the scalp along the sagittal sinus. The scalp skin was then removed, exposing a 5 mm × 5 mm area of the skull. Stainless steel screws were inserted into the cerebellum as ground electrodes. To expose the visual cortex, a craniotomy opening an area of 2 mm × 2 mm was performed followed by the removal of the dura mater. Implantation was conducted using a two-manipulator stereotaxic system. To secure the craniotomy, ceramic bone anchor screws and dental methacrylate were used to fix the flexible flat cable onto the mouse skull.

### Visual stimuli and electrophysiology recording

All visual stimuli were generated using MATLAB and the Psychophysics toolbox on a Windows PC. These stimuli were displayed monocularly on an ASUS PA248Q LCD monitor with a resolution of 1,920 × 1,200 pixels. The monitor was positioned 15 cm away from the eye. To ensure consistency in setup across experiments over time, physical bars were affixed to the air table as a reference to fix the monitor at the exact same location for each recording. We utilized a 30-second grayscale clip from the movie “Touch of Evil” (Orson Welles, Universal Pictures, 1958), which contained a continuous visual scene. To create the phase scrambled natural movies, we randomly scrambled the structure of the original clip, forming new versions of the movie^8^. In the case of the 50% and 100% phase scrambled movies, half and all of the image’s phase elements were randomized, respectively. The clip was contrast-normalized prior to the presentation and displayed at a frame rate of 30 frames per second. We conducted 20 repeats for each version of the movie. Each repetition commenced with a 5 s period of a gray screen, followed by the 30 s movie clip. For different mice, the cadence of the stimulation sessions is variable. The 9 mice were separated into two groups: frequent (average cadence < 8 days/stimulus session) and infrequent mice (average cadence > 15 days/stimulus session).

Electrophysiological recordings were performed throughout the entire course of the monthslong (∼4-6 months) visual stimulation experiment. Electrophysiological recordings were collected on a subset of the stimulus sessions, called recording sessions. For the same mouse, the neural signals were recorded consistently at either 10 kHz or 20 kHz with either a CerePlex Direct software (Blackrock Microsystems) or RHX Data Acquisition Software (Intan Technologies). Continuous signals were also notch filtered at 60 Hz (hardware filter) and bandpass filtered at 300–3,000 Hz using spikeinterface module in Python (https://spikeinterface.readthedocs.io/en/latest/) to subtract electrical artifacts associated with muscle activities such as those in the jaw and/or mystacial pad.

### Pupil tracking

To validate that the observed representational drift was not influenced by pupil dynamics, we conducted eye-tracking experiments involving four mice over ∼5 months. We recorded videos of the eye facing the LCD monitor by utilizing a customized infrared imaging system. To increase the contrast of the pupil, we employed an 850 nm infrared light source (LED Engin Inc). The eye and pupil dynamics were captured by a camera (35 fps). Subsequently, the frames in the videos were analyzed offline using DeepLabCut^43,44^ (https://github.com/DeepLabCut/DeepLabCut) to locate and track the eye and pupil throughout the video recording. Briefly, manually marking 16 points along the eye and pupil contours in representative frames (n = 150) generated a training set for a convolutional neural network (CNN) to learn the eye and pupil locations through transfer learning (ResNet-50). Remaining frames were then labeled by the trained CNN to label the corresponding marking points delineating the eye and pupil. Using Python scikit image (https://scikit-image.org/), the points were then fitted with two ellipses that estimate the eye and pupil on a frame-by-frame basis. The pupil-to-eye ratios were estimated by dividing the area of the fitted ellipse corresponding to the pupil by that corresponding to the eye on each frame.

### Spike sorting and curation

Common average reference was used to reduce the common-mode noise by creating an average of all electrode channels and subtracting it from each channel. For each mouse, the concatenated data from all recording sessions were then spike sorted using the fully automated algorithm Mountainsort 4.0^18^ (https://github.com/flatironinstitute/mountainsort) under the spikeinterface module; thus, the spike-sorting algorithm was blind to recording session and treated concatenated data as a single continuous signal. This eliminates the steps of matching units across multiple recording sessions. Spike threshold of >= 4σ was used for spike sorting. Signals from any electrode of interest and all five of its neighboring electrodes in individual hexode arrays were accounted for during spike sorting based on the designed device geometry.

The output templates of Mountainsort were first manually curated, during which we excluded from analysis any templates that corresponded to electrical noise based on waveform and mean waveforms plotted against the relative physical locations of the electrodes. Any unit templates whose interspike interval violation (< 1.5 ms) rate exceeded 10%, signaling great refractory period violation^15^, were also excluded. We further excluded single units that were not stable across multiple recording sessions. First, we compared individual single unit’s mean waveforms measured in different pairs of recording sessions by calculating the Pearson’s correlations. Those single units with high percentage of less correlated mean waveform pairs (> 10% of the correlations across different session pairs < 0.9) across sessions may be excluded after manual investigation, and those with highly similar mean waveforms will be treated as stable units and kept for downstream analysis. Then, the correlations of the mean waveforms across all pairs of units in the same hexodes were calculated to identify potential templates from the same units wrongly separated into two units. Those single-unit pairs with a great percentage of similar mean waveform pairs (> 90% of the across-session correlations between the two units > 0.8) may be merged after manual inspection. The individual units’ mean waveforms used in the above analyses are formed by concatenating its mean waveforms from all six neighboring electrodes in the corresponding hexode.

### Validation of stable units

#### Waveform stability

For all units, the within-unit correlations across consecutive recording sessions were compared with all across-unit waveform correlations within the same recording session. If any across-unit waveform correlation is higher than within-unit correlation, the unit will not be considered stable. Then, waveform features including amplitude, duration, repolarization slope, recovery slope, and peak-to-trough ratio were calculated using existing definitions^15^ for each spike of all stable units. The means of these waveform features in different sessions were then calculated and plotted for each stable unit to prove stable waveform over time. For all spikes in each stable unit, we then concatenated the waveforms from all six channels in its corresponding hexode to form concatenated waveforms. We then collected concatenated spike waveforms from all stable units on each hexode and applied uniform manifold approximation and projection^45^ (UMAP; https://umap-learn.readthedocs.io/en/latest/) to dimensionally reduce the spike waveforms to two dimensions (number of neighbors: 20; minimum distance: 0.1; metric of distance: Euclidean). Then, the two-dimensional points representing each spike’s concatenated waveform were plotted on a three-dimensional space for visualization, where the third dimension represents the recording session from which each waveform came. The locations and centroids of the clusters in different recording sessions were used to show long term stability of the waveforms of the stable units.

#### Estimation of position

The estimated position 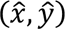 of source signals relative to the physical location of the electrodes on the hexode array is treated as the estimated centroid position of the single units, calculated with a previously defined method^6^:

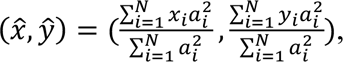

where a_i_ indicates the amplitude of the mean waveform collected on electrode i; x_i_ and y_i_ are the centroid coordinates of electrode i; and N is the number of electrodes. Note this is only an estimation of the unit centroid position instead of the actual location of the neuron. The estimated centroid position displacements of the same stable neuron between consecutive sessions were calculated and plotted (cumulative distribution and quintile lines) to prove minimal displacement (< 10 μm) of the stable units across consecutive sessions. The estimated centroid position distances between all pairs of different stable neurons on the same hexode in the same sessions were also calculated.

After proven stable with respect to waveform-defined stability and estimated centroid position-defined stability, the stable units are deemed as the same stably recorded neurons in the long term.

### Neuron-level analysis

From the spiking times of each stable neuron, mean firing rates were calculated for each time bin of either 0.25s or 0.5s throughout the 30-second movie by counting up the number of the spikes in single time bins and dividing it by the duration of the time bin. The firing rates of the same neurons in response to the same time bins (0.5 s) in the movie but in different recording sessions were then compared by computing Pearson’s correlation between responses from the previous sessions and those from latter sessions. A special case is comparing the response from the first and second ten repeats within the same sessions. In this analysis, we subtracted the mean baseline activities (when the movie is not playing) of each neuron from the neuron stimulus-dependent firing activities.

We then used existing definitions (including population vector, ensemble rate vector, and tuning curve vector)^7^ to characterize neuron-level representational drift. Time bin size is 0.5 s in these analyses. Briefly, the population vector (PV) contains the firing rates of the neurons in the population in response to any single time bin of the movie. Ensemble rate vector is the averaged firing rates of the neurons in the population over all PVs throughout the 30-second movie presentation. It is an indication of overall excitability of the neurons during each movie presentation. Tuning curve vector contains single neurons’ firing rates at all time bins during the 30 seconds of the movie. Next, for each of the three types of vectors, we calculated Pearson’s correlation within the session or across different pairs of sessions using a split-half approach. We split the 20 repeats into halves and averaged firing rates across the first 10 and latter 10 repeats of the session (producing matrices 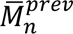 and 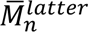 for session n) to compare within the same recording sessions. To compare across different pairs of recording sessions, we calculated the averaged matrices for the four halves 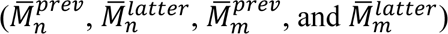 from the two sessions (m and n), which are then compared in a pairwise fashion. For PVs, only pairs of averaged PVs corresponding to the same time bins in the movie were correlated, and for tuning curve vectors, only pairs of averaged tuning curves for the same neurons were correlated. For each pair of halves, we then took the means of the PV correlations from all time bins and the means of the tuning curve vector correlations from all neurons. Last, the mean correlations (for PVs, ensemble rate vectors, and tuning curve vectors) averaged across different pairs of halves in all recording session pairs were correlated with the time interval between recording sessions. Additionally for PVs, we identified three consecutive recording sessions (3 days) in one mouse and calculated the mean firing rate matrices averaged across all 20 repeats 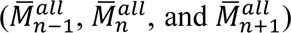. Then, we calculated the correlation of PVs between all pairs of different time bins from the three matrices. This allowed us to identify the internal structure^7^ of the neural response on each day and qualitatively observe its stability across consecutive days in the short term.

#### Experience number analysis

For all stimulus sessions throughout the multi-month experiment, session numbers were assigned to indicate the number of stimulus experiences the mice have received. Then, we used the previously calculated mean correlations of vectors (for all three types of vectors) between pairs of recording sessions and correlated them here with the difference in the number of experiences between pairs of recording sessions. This allows us to explore the effect of elapsed experience (instead of the elapsed time) on neuron-level representational drift.

#### Effect of time on drift

To measure the amount of drift from one recording session to the immediate next recording session, we calculated the mean cosine distance (CD) of the PVs averaged across all time bins between consecutive recording sessions, where cosine distance between two vectors x and y is defined as follows:

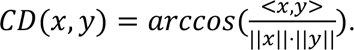

This follows the exact split-half method as before. To calculate the rate of drift^6^ from one recording session to the next, the cosine distance between all pairs of the halves from the two consecutive recording sessions (n and n+1) were first averaged 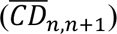 and then subtracted with a running average of the within-session PV cosine distance 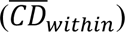, and finally divided by the amount of elapsed times in days (*T*_n+1_ − *T_n_*) between the consecutive recording sessions:

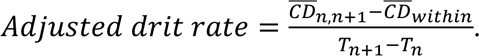

Next, we divide the monthslong experiment into two halves of equal time duration, each containing multiple recording sessions. Within each half of the recording sessions, we calculated the mean PV cosine distances averaged across all time bins and all pairs of the halves from the recording session pair. The mean cosine distances were then used to compute the drift rates between each pair of recording sessions. Since the analysis is not between consecutive sessions but rather between all possible session pairs, there is no running average of within-session PV cosine distance to subtract from the mean cosine distance. The drift rate is then defined as follows:

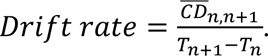

This allowed us to compare the drift rate of PVs in the previous and latter halves of recording sessions during the monthslong stimulation experiment.

#### Visual response geometry

Visual response geometry is defined by the geometry formed by high-dimensional points representing PVs in response to all time bins (0.5 s) during the movie in the space whose dimensions represent the firing rates of individual neurons in the population. For each recording session, the points represent the mean PVs averaged across all 20 repeats. To characterize the constancy/inconstancy of the visual response geometry, all angles formed by all sets of three high-dimensional points were calculated and compared across pairs of recording sessions, based on an existing method^6^. Briefly, all angles from one recording session formed an edge matrix for that session. The Frobenius norms of the difference between pairs of edge matrices from different pairs of recording sessions (session m and n), 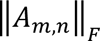, were computed and rescaled using the following approach. The edge matrices in the first and last recording sessions were shuffled along the dimension of time bins, and the mean Frobenius norm of their difference, 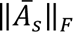, was calculated. Then, the edge matrices from the first and latter halves of the 20 movie repeats were calculated, and the Frobenius norm of their difference was calculated for each recording session to yield the within-session Frobenius norm. The mean of the within-session Frobenius norm, 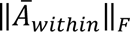, was calculated across all recording sessions. The across-session Frobenius norm is then rescaled:

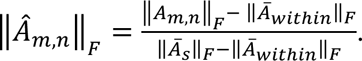

#### Parametrically scrambled movies

For each of the differentially scrambled movies, the tuning curve vectors were compared across all the halves from pairs of recording sessions using Pearson’s correlation and cosine distance. The mean correlations of the tuning curve vectors averaged across different pairs of halves in different pairs of recording sessions were then computed and plotted against the time interval between the recording sessions. Note, this tuning curve vector analysis for the differentially scrambled movies employed the same method as that for the original movie. Next, the drift rate of the tuning curve was calculated from the difference in cosine distance using aforementioned methods (not adjusted version). The turning curve vector correlation and drift rates were then pooled across all time intervals and compared across the three differentially scrambled movies.

### Population level analysis

Gaussian process factor analysis (GPFA)^24^ was used to expose the low-dimensional neural latent dynamics in response to the natural movie stimuli. For each mouse, the neural firing rate data (0.5-second bin) from all recording sessions (each with 20 movie repeats) were concatenated, and a single GPFA was applied to reduce the dimension from the number of all stable neurons in each mouse to five. Concatenating the data from multiple recording sessions and applying one single GPFA can produce a single linear neural manifold/subspace on which latent dynamics from different recording sessions reside. For each of the five dimensions, the projection of the latent dynamics from different pairs of recording sessions were compared. Specifically, the projections of 20 repeats were averaged for each recording session, and the averaged projections were then compared using either Pearson’s correlation or cosine similarity between different pairs of recording sessions. The cosine similarity and Pearson’s correlation between two vectors x and y are defined as follows:

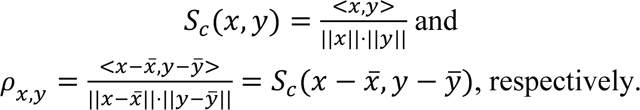

Note that Pearson’s correlation is the same as the cosine similarity if the vectors are centered by the mean. The average Pearson correlation or cosine similarity was then calculated across the different pairs of the recording sessions. This allows us to select the three dimensions with the top correlation or cosine similarity, forming the three-dimensional latent dynamics for downstream analysis and visualization. Note, however, the latent dimensions are not numbered based on the values of correlation or cosine similarity.

To further compare the latent dynamics and investigate their correlation/similarity at various time intervals, the 20 repeats were split into two halves in each recording session. For each half, we then averaged the latent dynamics from all 10 repeats to get two averaged latent dynamics in one recording session. Next, for each dimension, a Gaussian filter (σ=3) was applied to generate two smoothened latent dynamics for each recording session. To compare the stability of latent dynamics within the same sessions, the smoothened latent dynamics on each dimension from the two halves were compared and averaged. To compare the stability of latent dynamics across different sessions, the smoothed latent dynamics on each dimension from each half in one recording session were compared with that from each half in the other session, forming a total of 4 comparisons for each pair of recording sessions. The means of the correlations or cosine similarity from the 4 comparisons were taken. The mean correlation or cosine similarity from all three dimensions were then taken as an indicator of the overall correlation or similarity of the latent dynamics at the two times. Finally, the correlations or cosine similarity between the latent dynamics from pairs of recording sessions were then correlated with the time intervals between pairs of recording sessions.

For the two differentially scrambled movies (50% and 100%), we also applied a single GPFA to the neural data combined from all recording sessions. The means of the correlations across the three dimensions from different pairs of recording sessions were compared and plotted against the time intervals, following previously described methods.

#### Transformation of latent dynamics

To attempt to recover more correlated and similar latent dynamics over the long term, we separately applied GPFA to the neural activity data in each recording session, which serves to translate the neural data by centering it. This also produces the latent dynamics and the neural manifold defined by the loading matrix given by GPFA. In order to compare the latent dynamics, the neural manifolds between consecutive sessions are aligned using a previously defined approach^11^. Specifically, after dimensionally reducing the neural data (0.5-second time bins) in each session from the number of all neurons to 5, the corresponding loading matrix (Λ), defining the neural manifold, in the next session will be multiplied by an orthogonal matrix O so that it can match maximally the loading matrix in the previous session.

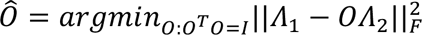

This is referred to as the orthogonal Procrustes problem, which has a closed form solution^46^. This way, the loading matrices and thus the neural manifolds between consecutive recording sessions are maximally similar. This stage is referred to as the “translated” stage. To compare the latent dynamics, the top three dimensions with the highest correlations or cosine similarity were selected, and the means of the correlations or cosine similarity across the three dimensions were taken as the similarity of the latent dynamics from each pair of recording sessions, following the aforementioned approaches.

To then account for any transformation of the latent dynamics across consecutive recording sessions, we then attempted to apply one linear transformation (A) to vectors of the time-varying latent dynamics (z) at each time bin so that the latent dynamics in the next recording session can maximally match that in the previous session.

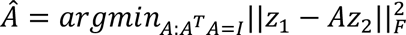

This forms another orthogonal Procrustes problem. Applying a linear transformation to the latent dynamics is equivalent to applying a linear transformation to the loading matrix, thus changing the neural modes. This stage is referred to as the “transformed” stage. All five dimensions were used in this step. To compare the transformed latent dynamics using either correlation or cosine similarity, the same aforementioned approaches were used, during which the same three dimensions from the “translated” stage were used.

#### Simulation of probe instability

To compare the ability to extract highly correlated and similar latent dynamics from stable and unstable neural recordings, we simulated two modes of probe instability: unit change from drifting probe and unit dropout from broken channels^11^. Specifically, probe drift is equivalent to the micromotion between the neurons and the electrode, leading to a change of the recorded neurons. Therefore, this is simulated by holding out a subset (30% of the original neuron population) of neurons and randomly substituting the activities from a percentage of remaining neurons with those in the hold-out group. For unit drop out, the selected remaining neurons’ activities were substituted with an activity pattern sampled from a standard normal distribution mimicking noise.

Different levels of probe instability were simulated: 0%, 10%, 20%, and 30%. The percentage of the probe instability represented the percentage of unit dropout as well as the percentage of the unit change in the simulated final recording sessions. Specifically, the unit dropout and unit change were to happen gradually in randomly sampled recording sessions, and the percentage represents the level of probe instability in the last recording session. The units to be dropped out and those to be changed were mutually exclusive. Two mice from each stimulus frequency group were selected for this analysis. For each level of probe instability, 10 simulations were performed. For each simulation, the simulated data from each recording session was separately applied with GPFA for dimension reduction. Later, the neural manifolds were aligned, and latent dynamics were translated and transformed across consecutive recording sessions. The transformed latent dynamics were then compared using correlation with the aforementioned methods.

### Machine learning based decoders

Machine learning and artificial neural networks (ANN) were used to evaluate the encoding of the visual information in the neural data. Specifically, the neural data (0.25-second time bin) throughout the 30-second movie were divided into 8 time windows, each with 15 time bins. Then, the neural data in each time window and the corresponding window number form input and output pairs for training and evaluating a recurrent ANN (RNN) with the long short-term memory (LSTM) architecture and a support vector machine classifier model. The two models are intended to predict the identity of the time window after taking in the neural data from arbitrary time windows as input. To evaluate the same model’s ability to predict the time window throughout the monthslong recording period, we divide the recording sessions of each mouse into two halves, each spanning multiple months. We then used the data in the first half to train the model (cross entropy loss and Adam optimization) and used the data from the latter half for evaluation. During training, 10% of the data were left out for validation and model selection, and for the remaining 90% of the data, 5-fold cross validation in combination with the grid search was used to find the set of parameters that generates the minimal mean loss across the 5 folds.

Three input data types were used in LSTM and SVM models. First, the original high-dimensional neural data containing the firing rates of each individual neuron was used. Two types of low dimensional neural data at different stages were also used. Specifically, one GPFA was applied to the combined neural data from all recording sessions and to reduce the dimensions. The dimension was reduced from the total number of neurons to a level with no less than 95% of the explained variance. The cumulative values from the top D eigenvalues out of the sum of all eigenvalues in the covariance matrix of the centered trial-combined data^47^ is used to inform the explained variance of top D dimensions. The low-dimensional data at this stage (pre-translation) was used as the second type of input. Then, separate GPFA were applied to the neural data from each recording session and to reduce the dimensions to the same level as before, which translated the neural dynamics. Then the manifolds across consecutive recording sessions were aligned, and the latent dynamics were transformed. This produced the third type of input: a low-dimensional latent dynamics at the “transformed” phase.

The LSTM framework in PyTorch (https://pytorch.org/) in Python was used. During training and evaluation, the time bins in each time window were sequentially passed into the RNN model with two layers. During grid search, combinations of the following parameters were explored: dimension of the hidden layer (25 and 50), drop out ratio (0.2 and 0.3), learning rate (0.001 and 0.005), and batch size (50 and 100). The outputs from two LSTM layers were then concatenated and passed into one densely connected linear layer with the leaky rectified linear unit (Leaky ReLU) as nonlinear activation function. In the output layer, softmax function was used to make a prediction about the identity of the time window. The SVM classifier implemented in scikit-learn (https://scikit-learn.org/stable/) in Python was used. During training and evaluating, the time bins in each time window were concatenated and passed into the model as a whole. During grid search, various combinations of the regularization parameter, C (0.0001, 0.0005, 0.001, 0.005, 0.01, 0.05, 0.1, 0.5, 1, 5, 10, and 50), and kernel types (linear, polynomial, radial basis function, and sigmoid) were explored. Analyses of both types of models were applied for each individual mouse and each type of input data.

### Statistics

Statistical analysis was performed using GraphPad Prism 10. The data first underwent a normality test. If the data passed the normality test (*P* < 0.05), parametric analysis was performed. If not, nonparametric analysis was performed. Then, if the data can be paired, paired statistical tests were used. Otherwise, unpaired tests were chosen. All statistical tests were two-tailed if applicable and used a *P* value of 0.05 to trigger significance. Notably, when comparing the drift rates of tuning curve vectors among differentially scrambled movies, any drift rates smaller than 0 (which was possible under the calculation of adjusted drift rate) were excluded, leading to the use of unpaired test: ordinary one-way analysis of variance.

For all the linear regressions, linear regression models with ordinary least squares under the statsmodels API module (https://statsmodels.org) in Python were used. Pearson’s correlation coefficients and the *P* value were reported for all linear regressions. Bootstrapping was conducted 1000 times, and linear regression was fitted to each resampled data. At each value along the abscissa, the 2.5 percentile and 97.5 percentile of the values generated by the 1000 regression lines were taken as the 95% confidence intervals in the plots.

## Data availability

The source data supporting the main results of this study are available with the paper.

## Acknowledgement

We acknowledge the discussion and assistance from all Liu Group members. We acknowledge the support from the Harvard University School of Engineering and Applied Sciences Startup fund and the Harvard University Faculty of Arts and Sciences Dean’s Competitive Fund for Promising Scholarship. The funders had no role in study design, data collection and analysis, decision to publish or preparation of the manuscript. We thank scidraw.io for illustrations.

## Author contributions

J. Liu, S.Z., and H.S. conceived and designed the experiments. S.Z. and J. Lee fabricated and characterized the electrodes. S.Z. and S.J. performed the brain implantation and in vivo recording. H.S., M. L., and S.Z. conducted the data analysis. J. Liu, H.S., and S.Z. wrote the manuscript. All authors discussed and commented on the manuscript.

## Extended Data Figures and Extended Data Figure Captions

**Extended Data Fig. 1.**
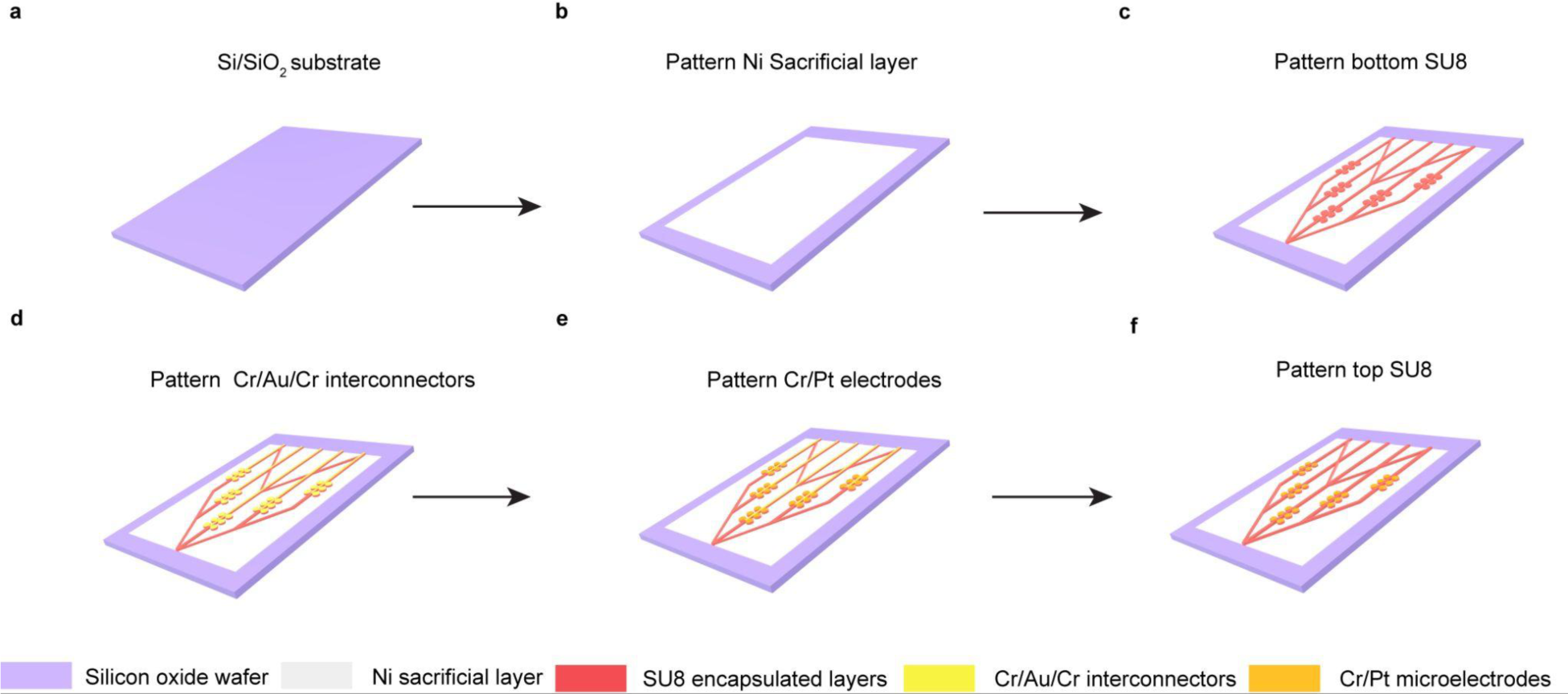
Fabrication steps of tissue-like flexible mesh electronics with hexode electrode arrays. **a,** The 3-inch Si/SiO_2_ wafers were used as insulating substrates for the fabrication process of hexode mesh electronics. **b,** Ni sacrificial layers were defined by photolithography and deposited through thermal evaporation. **c,** SU-8 2000.5 bottom passivation layer (red) was defined by photolithography on the Ni sacrificial layer. **d**, The Cr/Au/Cr interconnects were defined by photolithography and deposited through electron beam (e-beam) evaporation on top of the SU-8 passivation layer. **e,** Cr/Pt microelectrodes were defined by photolithography and deposited through e-beam evaporation. **f,** SU-8 2000.5 top passivation layer (red) was defined by photolithography.

**Extended Data Fig. 2.**
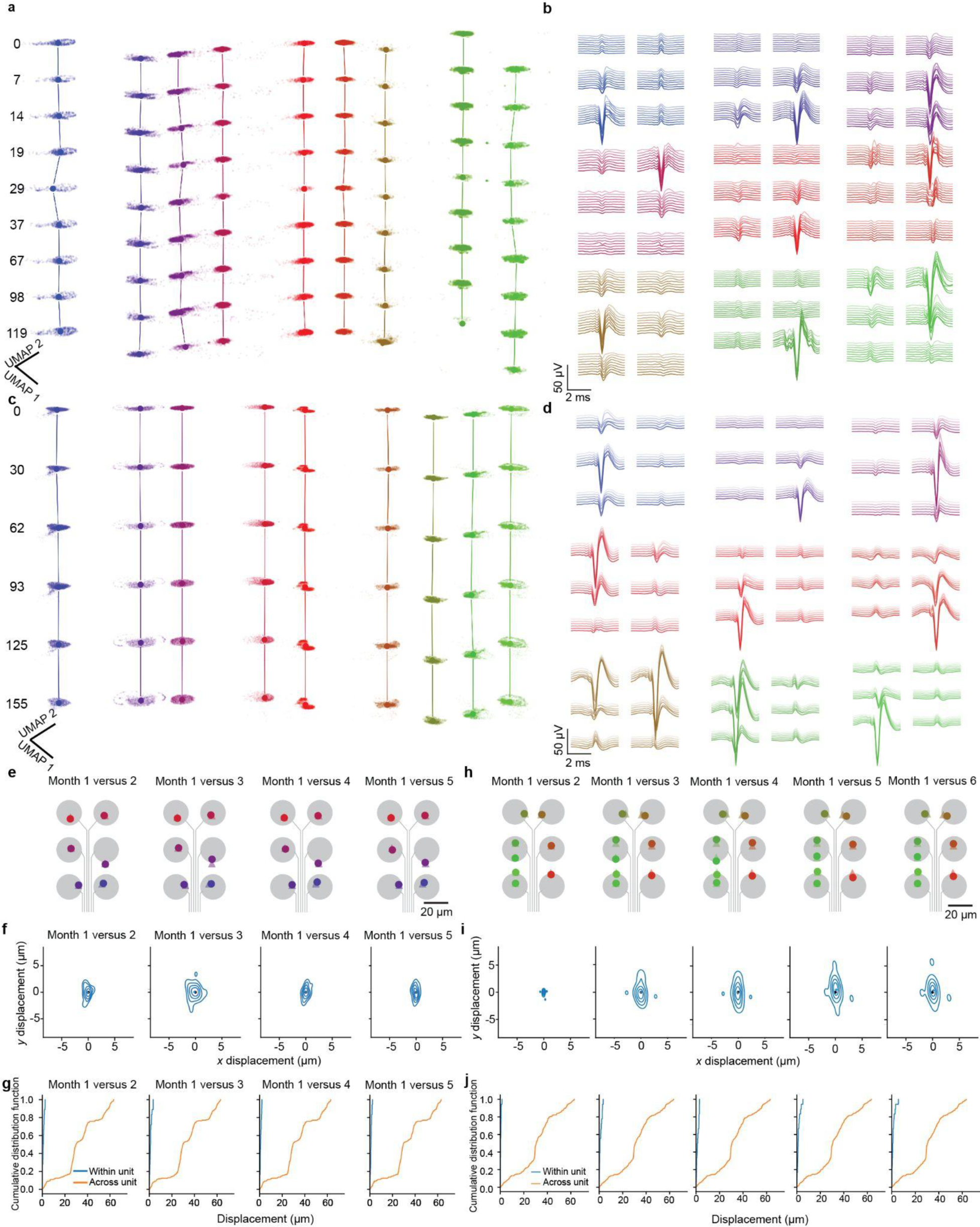
Long-term stable tracking of the same-neuron activities over multiple months reflected by waveform-defined and neuron-centroid-position-defined similarity. **a,** Uniform manifold approximation projection (UMAP) of each spike’s waveforms from 9 representative neurons from all recording sessions (n = 9) in a mouse with frequent visual stimulation. Each cluster is composed of points representing a waveform embedded on a 2-dimensional UMAP embedding space. The z-axis represents the different recording sessions over multiple months with the days indicated. **b,** Representative average waveforms of neurons in **a** recorded simultaneously at each of the six recording electrodes of hexode mesh electronics. Waveforms recorded at different times are plotted in order of gradient color. **c,** Same as in **a** but for a mouse with infrequent visual stimulation. **d,** Same as in **b** but for the mouse in **c**. **e,** Estimated single-neuron centroid positions on one representative shank of the mesh electronics throughout the 5-month recording from the mouse in **a**. The estimated centroid position for each single neuron on month 1 (triangle) was compared to those on months 2-5 (circle). Gray circles indicate the positions of the individual electrodes within an array. **f,** Mean displacement of single-neuron estimated centroid position from the mouse in **e** between month 1 (black dots, defined at origin) and months 2-5 (blue dots; columns 1–4, respectively; not visible due to overlapping with black dots). Contours indicate quintiles of the distribution of estimated centroid position displacement for the neurons. **g**, Cumulative distribution of within-neuron estimated centroid position displacement (blue) between month 1 and months 2-5 (columns 1–4, respectively) and across-neuron estimated location centroid distance within each recording session (orange) for the mouse in **e**. **h-j,** same as in **e-g** but for the mouse in **c-d.**

**Extended Data Fig. 3.**
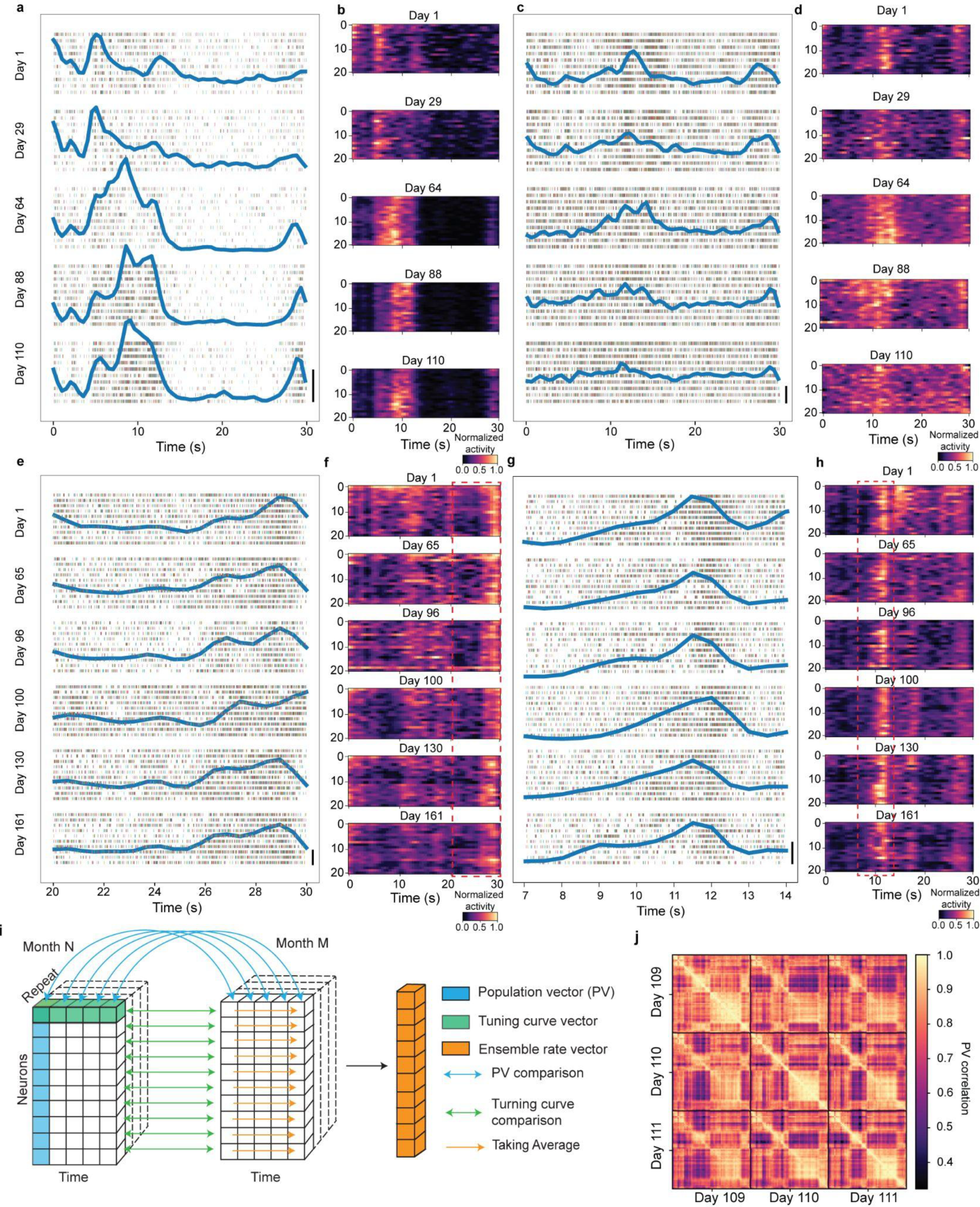
Visual stimulus-dependent neural activities in the mouse primary visual cortex and relevant metric definitions. **a-d,** The responses of two representative neurons from two mice with frequent visual stimulation. In **a** and **c**, single spikes of two neurons (from two mice) in response to 10 repeats (odd repeats out of the 20 repeats) of the movies in representative sessions are shown in the raster plots, and the normalized all-repeat-averaged peristimulus time histogram (blue lines) are plotted on top of the raster plots. Scale bars correspond to a value of 0.2 AU of the normalized firing rate. The smoothed firing rates of the two neurons for all repeats (n = 20) in representative sessions are plotted in **b** and **d**, respectively. **e-h,** The responses of two representative neurons from two mice with infrequent visual stimulation. In **e** and **g,** single spikes of two neurons from two mice in response to 10 repeats (odd repeats out of the 20 repeats) of the movies in representative sessions are shown in the raster plots. The time windows for the two representative mice are 10 seconds and 7 seconds, respectively. This time duration corresponds to the red box in **f** and **h**. The normalized all-repeat-averaged peristimulus time histogram (blue lines) for corresponding time windows are plotted on top of the raster plots. Scale bars correspond to a value of 0.2 AU of the normalized firing rate. The smoothed firing rates of the two neurons for all repeats (n = 20) of the entire 30-second movie in representative sessions are plotted in **f** and **h**, respectively. **i**, Schematics showing data structure, definitions of population vector (PV), ensemble rate vector, and tuning curve vector, and the calculation of the correlations or cosine similarities. **j**, The correlations between population vectors from all time points in the movie from three consecutive recording sessions (from three consecutive days). Each block represents the population vector correlations of all time points in the movie from two recording sessions.

**Extended Data Fig. 4.**
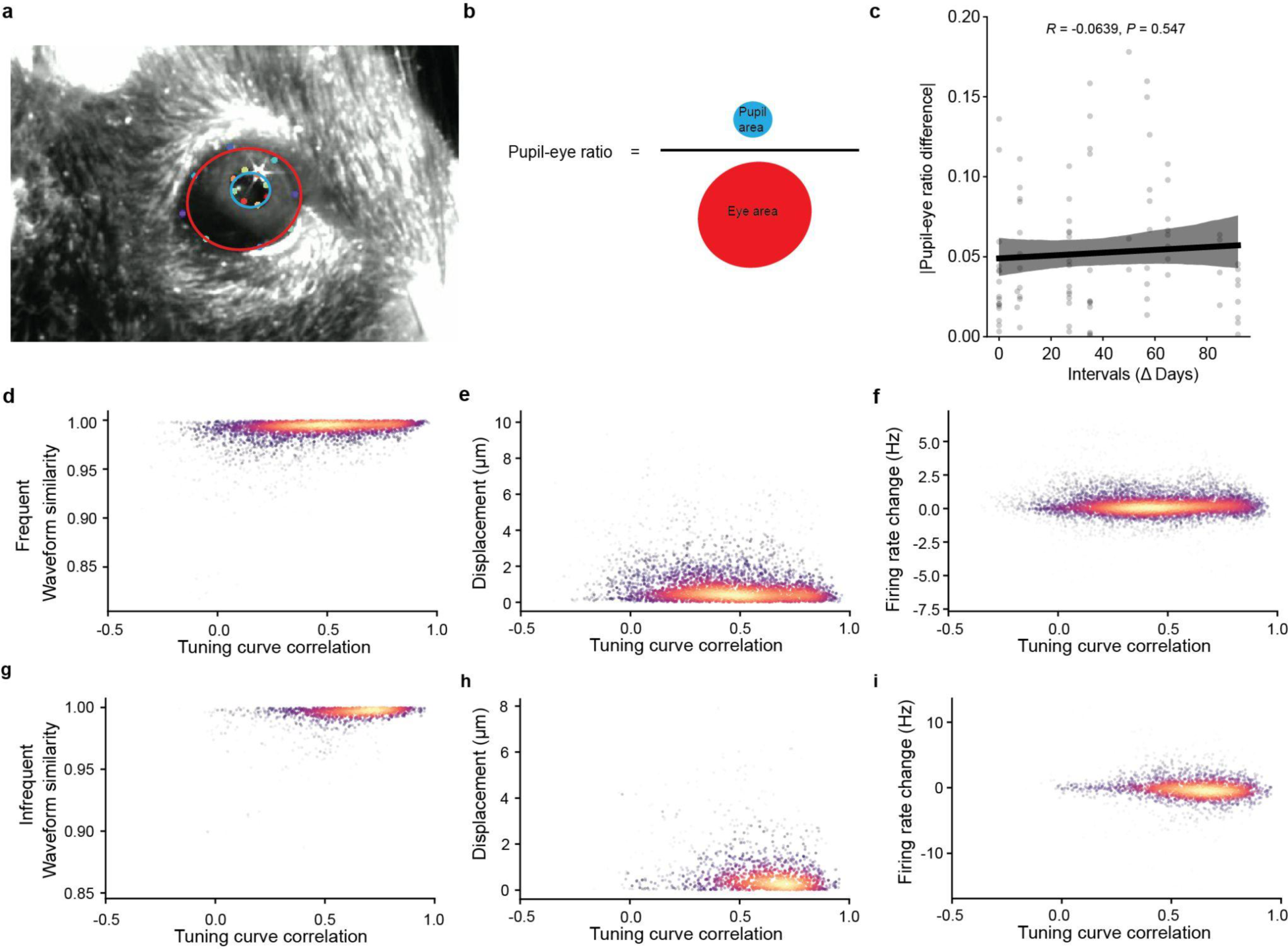
Independence of changes in stimulus-dependent single-neuron firing dynamics on potential confounding factors. **a**, Photograph showing a single frame of a mouse eye. The pupil and eye were delineated with dots using deep learning approaches and fitted with ellipses (red for eye, blue for pupil). **b**, Definition of pupil-to-eye ratios. **c**, Changes in mean pupil-to-eye ratios plotted against time intervals. The straight line with shaded area is the regression line with 95% CI of the linear regression. **d-f,** Scatter plots color-coded by density demonstrating high waveform similarity (**d**), low estimated neuron centroid position displacement (**e**), and near-zero firing rate change (**f**) with varying tuning curve correlations for single neurons (n = 84 neurons) in four independent mice stimulated frequently. **g-i,** Same as **d-f** but for five independent mice with infrequent stimulation (n = 130 neurons).

**Extended Data Fig. 5.**
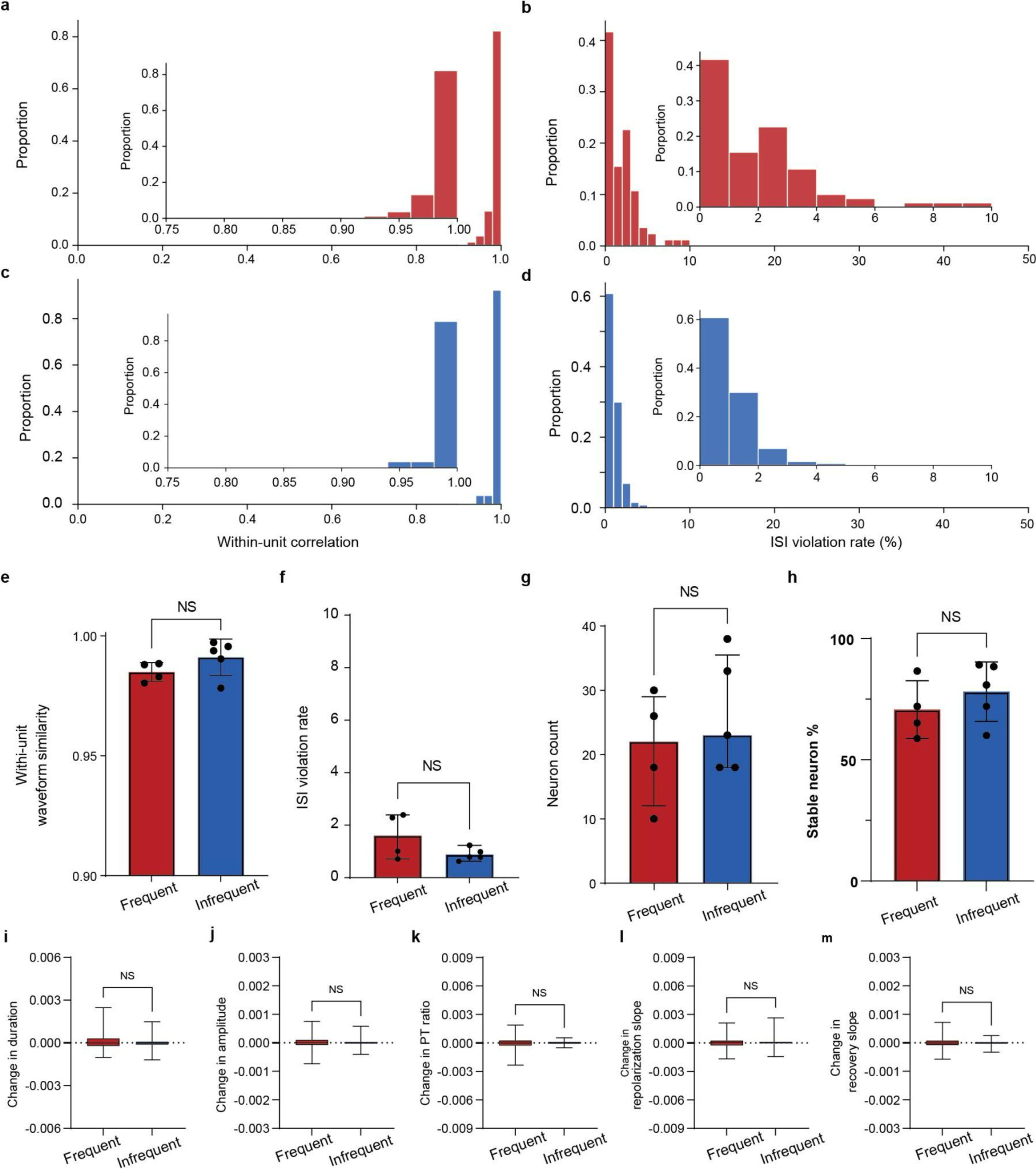
Comparing the statistics of neurons and waveforms in both groups of mice with frequent and infrequent stimulation. **a-b,** Histograms of within-neuron waveform correlations (**a**; bin size = 0.02) and interspike interval (ISI) violation rates (**b**; bin size = 1%) of the neurons (n = 84 neurons) in four independent mice with frequent stimulations. Inset is the zoomed-in view of the histogram. **c-d**, Same as in **a-b** but for five independent mice with infrequent stimulations (n = 130 neurons). **e-h**, Comparison of within-neuron waveform correlations (**e**; *P* = 0.1907), ISI violation rates (**f**; *P* = 0.1138), total neuron counts (**g**; *P* = 0.4345), and stable neuron percentage out of all sorted units (**h**; *P* = 0.3921) between the four independent mice with frequent stimulation and five independent mice with infrequent stimulation. Two-tailed unpaired t-tests were performed for all four pairs of comparisons. **i-m,** The rate of change of neuron waveform features - duration (**i**; *P* = 0.4313), amplitude (**j**; *P* = 0.2021), peak-trough ratio (**k**; *P* = 0.9062), repolarization slope (**l**; *P* = 0.5249), and recovery slope (**m**; *P* = 0.8643) - with respect to the time course during the multi-month recording for the mice with frequent stimulation and those with infrequent stimulation. Two-tailed Mann-Whitney tests were performed for all five pairs of comparisons.

**Extended Data Fig. 6.**
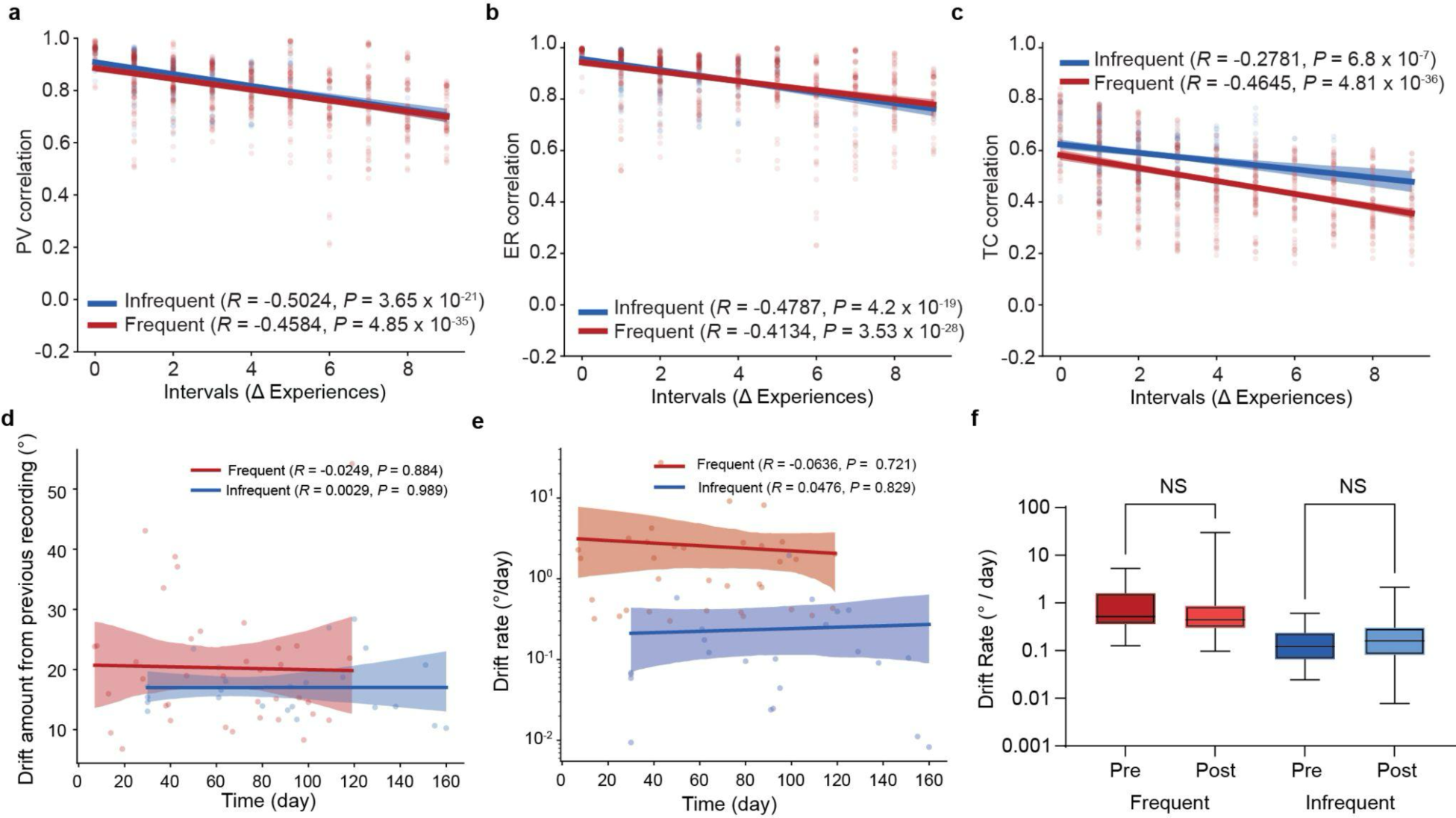
Effect of experiential number difference and time on representational drift in mice with frequent and infrequent stimulation. **a-c,** The correlations of PVs (**a**), ensemble rate vectors (**b**), and tuning curve vectors (**c**) between a subset of the recording session pairs plotted against the corresponding experience number difference for both mice with frequent (*n* = 4 mice) and infrequent (*n* = 5 mice) visual stimulation. The subset of recording session pairs includes all session pairs with an experience difference smaller or equal to 9. **d,** The amount of drift measured by the change in cosine distance between population vectors across consecutive recording sessions. Each point represents the drift amount from the previous recording session to the current recording session. **e,** The rate of drift at different times. Each point represents the drift rate calculated from the previous recording session to the current recording session. **f,** Comparison of the pooled drift rates from session pairs of all time intervals from the previous and latter half of the multi-month recording period in both mice with frequent and infrequent visual stimulations (red boxes, *P* = 0.3993; blue boxes, *P* > 0.9999; Kruskal-Wallis test with Dunn’s multiple comparison test).

**Extended Data Fig. 7.**
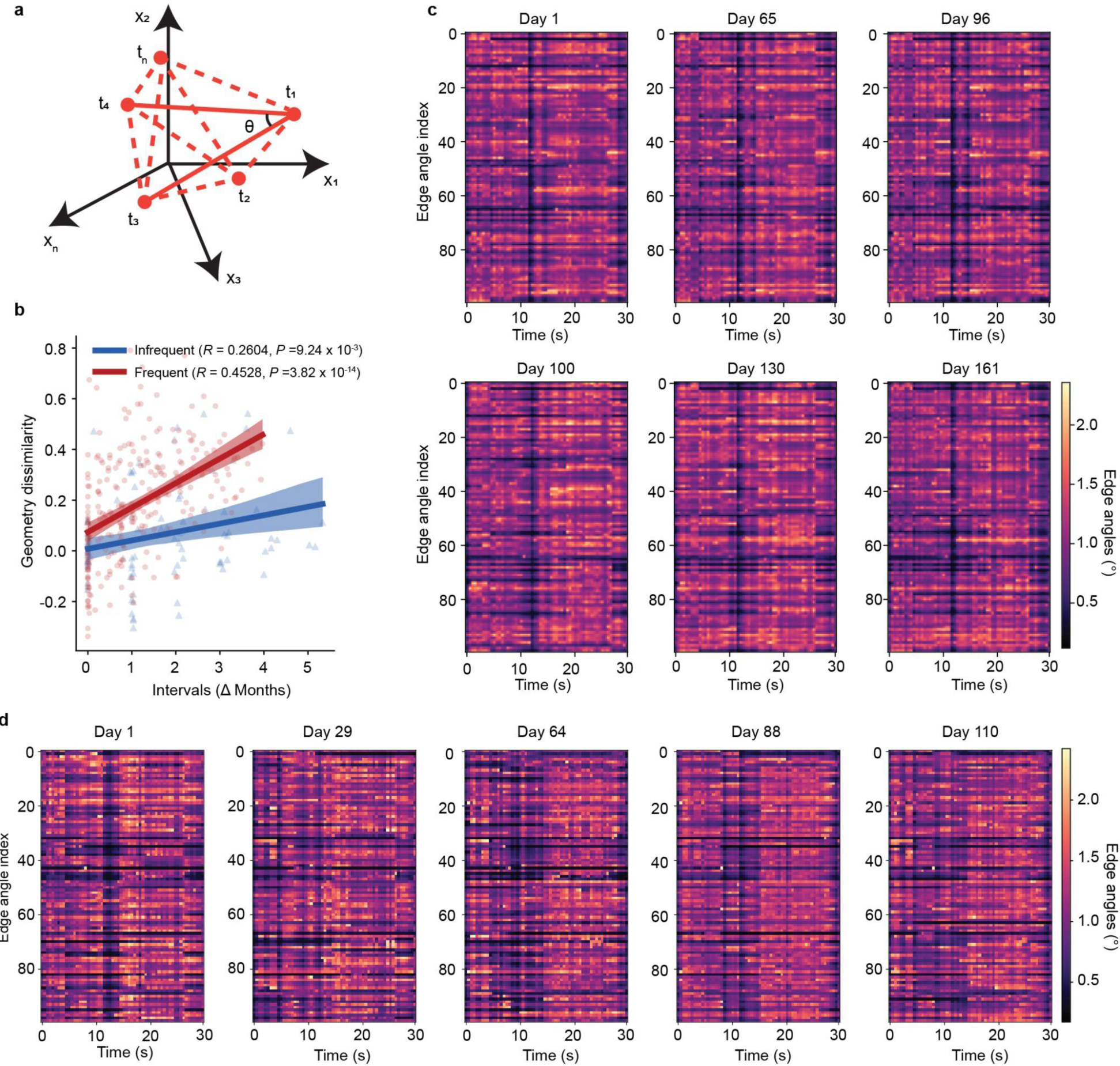
Change in the edge angles of visual response geometry over multiple months. **a,** Schematics of the visual response geometry. Each point represents the responses of all neurons in the population to a specific time bin in the 30-second movie. Each axis corresponds to one neuron’s responses. All the edge angles, which are formed by all sets of three points in the visual response geometry, can be arranged into a matrix, whose long-term stability is quantified using the Frobenius norm. **b**, The dissimilarity in the angles between pairs of recording sessions plotted against the time intervals, for both mice with frequent and infrequent stimulations. **c-d,** The matrices of the same set of randomly sampled edge angles in different recording sessions for two representative mice with infrequent (**c**) and frequent (**d**) stimulation, respectively.

**Extended Data Fig. 8.**
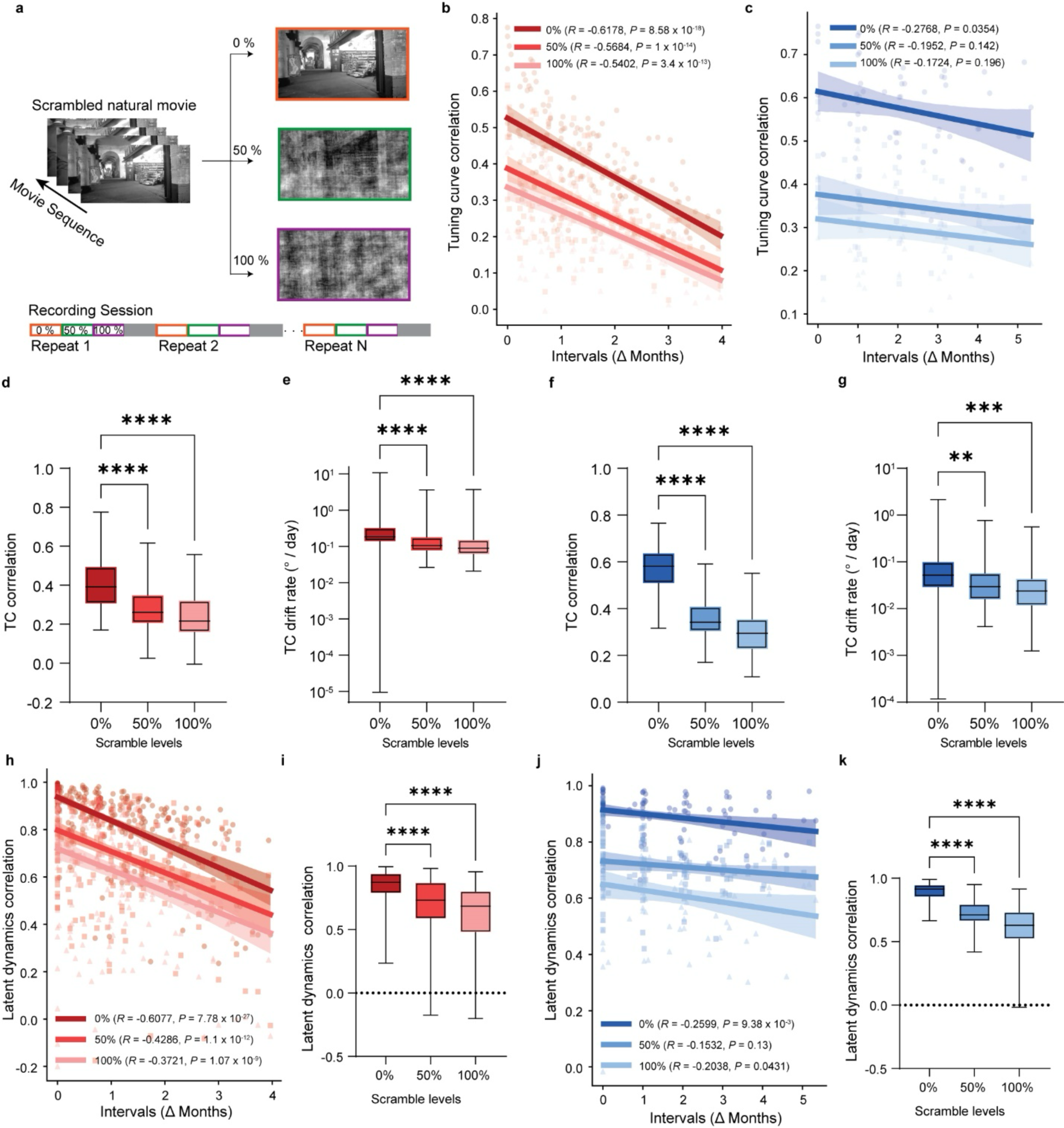
High-order statistics of visual stimulus affecting single-neuron-level firing dynamics and population-level latent dynamics. **a,** Parametric phase scrambling of the movie into three levels: 0% (original movie), 50%, 100% (noisy pixels). The movie at each level was presented to the mouse for 20 repeats. **b-c,** The correlation between tuning curve vectors from pairs of recording sessions in response to the movies at all three levels of parametric scrambling plotted against the time intervals, for both mice with frequent (**b**) and infrequent stimulations (**c**). **d-e,** The tuning curve correlation (**d**; *P* < 0.0001 for both comparisons; Friedman test and Dunn’s multiple comparison test) and drift rates (**e;** *P* < 0.0001 for both comparisons; Kruskal-Wallis test and Dunn’s multiple comparison test) pooled from all time intervals in all mice with frequent stimulation. Cosine distance was used to calculate the rate of drift of tuning curve vectors. The bars are plotted with 95% confidence intervals. **f-g,** Same as **d-e** but for mice with infrequent stimulation (**f**, *P* < 0.0001 for both comparisons; repeated measure one-way ANOVA with Geisser-Greenhouse correction and Dunnett’s multiple comparison test; **g,** *P* = 0.0043 and 0.0002 for 0% vs 50% and 0% vs 100% scrambling level, respectively; Kruskal-Wallis test with Dunn’s multiple comparison test). **h,** The averaged correlations (across the top three latent dimensions) between latent dynamics from pairs of recording sessions in response to the movies at all three levels of parametric scrambling plotted against the time intervals, for mice with frequent stimulation. **i,** Comparing the averaged correlations of latent dynamics pooled from all session pairs in **h** in response to movies at the three levels of parametric scrambling (*P* < 0.0001 for both comparisons; Friedman test and Dunn’s multiple comparison test). **j,** Same as in **h**, but for mice with infrequent stimulation. **k,** same as in **i** but for data in **j** (*P* < 0.0001 for both comparisons; Friedman test and Dunn’s multiple comparison test).

**Extended Data Fig. 9.**
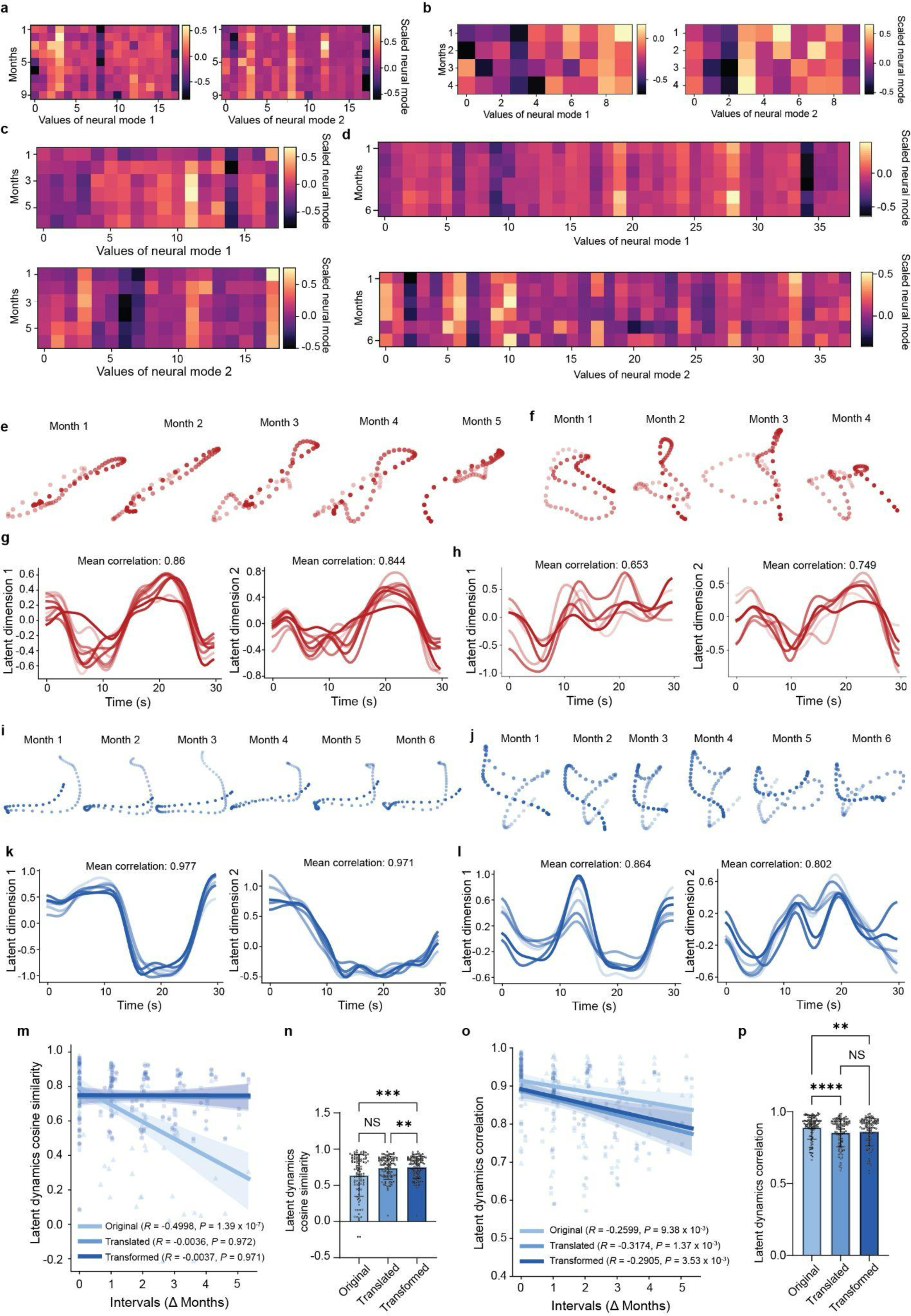
Latent dynamics of neuron population after translation, manifold alignment, and transformation for both groups of mice with frequent and infrequent stimulations. **a-b,** Neural modes (vectors defining each axis of the neural manifold) in different recording sessions for the first (left) and second (right) latent dimensions after latent dynamics translation for two mice with frequent visual stimulation. Each row in the blocks represents a vector representing a neural mode. **c-d,** Same as **a-b** but for two mice with infrequent visual stimulation. First and second latent dimensions are plotted on top and bottom, respectively. **e-f,** Latent dynamics after transformation for the two mice (**e** and **f**) respectively corresponding to the mice in **a** and **b**. **g-h,** Latent dynamics after transformation projected onto the first (left) and second (right) latent dimensions for the two mice (**g** and **h**), respectively corresponding to the mice in **a** and **b**. **i-j,** Latent dynamics after transformation for the two mice (**i** and **j**) respectively corresponding to the mice in **c** and **d**. **k-l,** Latent dynamics after transformation projected onto the first (left) and second (right) latent dimensions for the two mice (**k** and **l**), respectively corresponding to the mice in **c** and **d**. **m**, Averaged cosine similarities of latent dynamics between pairs of recording sessions before (original, same as in Fig. 4) and after latent dynamics translation and manifold alignment (translated), and after transformation (transformed), plotted against different time intervals for mice stimulated infrequently. **n**, Bar plot showing the mean, standard deviation, and distribution of the cosine similarities of latent dynamics between all pairs of sessions at the three stages of transformation (original vs translated: *P* > 0.9999; original vs transformed: *P* = 0.0002; translated vs transformed: *P* = 0.0013; Friedman test and Dunn’s multiple comparison test). **o**, Same as in **m** but for the averaged correlations. **p**, Same as in **n** but for the averaged correlations (original vs translated: *P* < 0.0001; original vs transformed: *P* = 0.0076; translated vs transformed: *P* = 0.0906; Friedman test and Dunn’s multiple comparison test).

**Extended Data Fig. 10.**
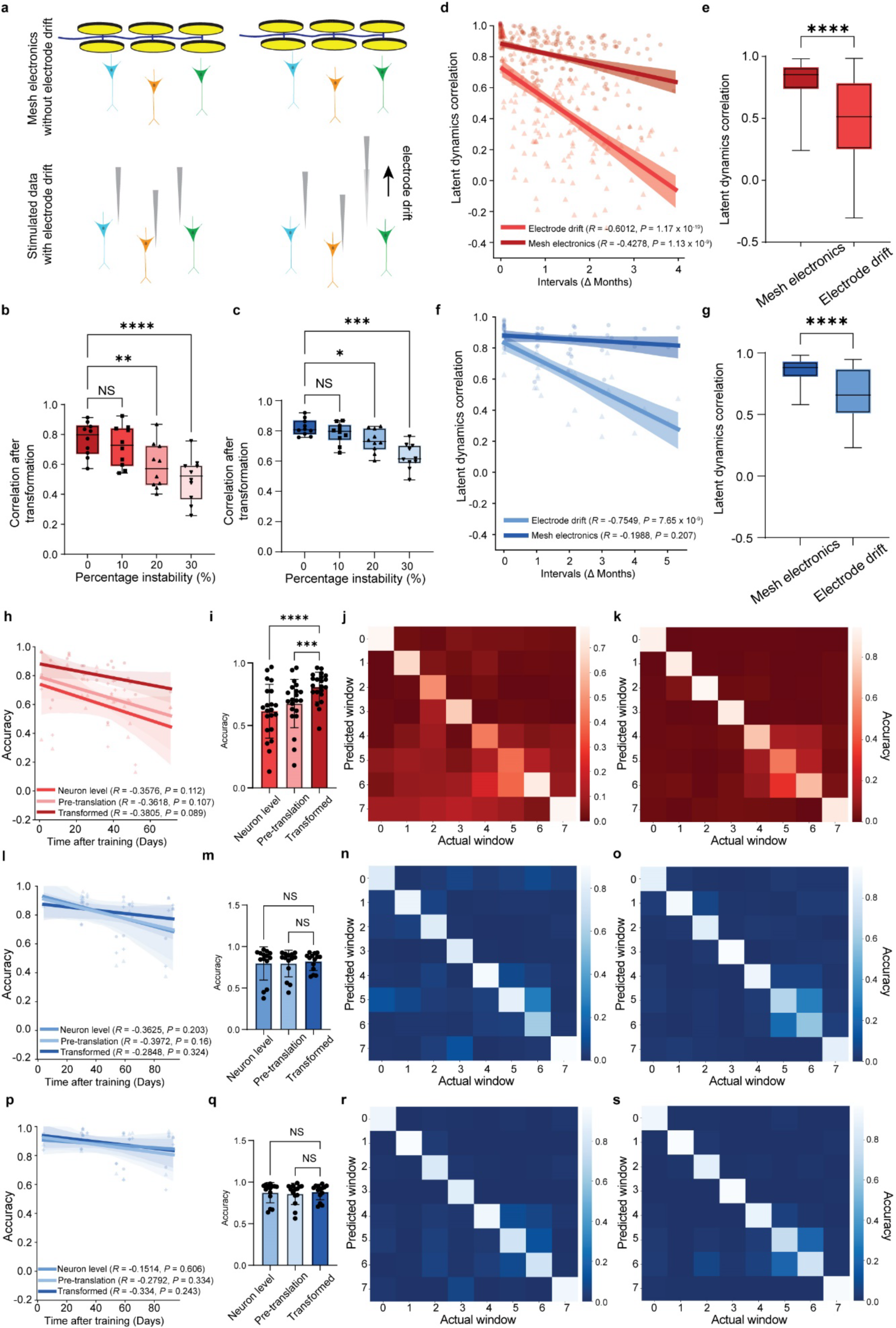
Flexible mesh electronics enabling high performance in latent dynamics transformation and visual information decoding in V1 with representational drift. **a,** Stability of mesh electronics, which has minimal probe drift, in tracking the same neurons (left) and common modes of probe instability such as probe drift (right). **b,** Means of averaged correlations (across the top three latent dimensions) between latent dynamics in different session pairs after transformation of latent dynamics, calculated from simulated data with electrode drift at different percentages (10%, 20%, and 30% of neuron tuning change and neuron dropout in the last session). Each point represents the mean of the averaged latent dynamics correlations from one simulation of latent dynamics transformation (a total of 10 simulations from two representative mice with frequent stimulations; 5 simulations for each mouse). *P* = 0.5547, 0.0012, and < 0.0001 for no electrode drift vs 10%, 20%, and 30% of electrode drift, respectively (repeated measure one-way ANOVA with Geisser-Greenhouse correction and Dunnett’s multiple comparison test). **c**, Same as in **b**, but for mice with infrequent stimulations. *P* = 0.2095, 0.0125, and 0.0001 for no electrode drift vs 10%, 20%, and 30% of electrode drift, respectively (repeated measure one-way ANOVA with Geisser-Greenhouse correction and Dunnett’s multiple comparison test). **d,** All averaged correlations (across the top three latent dimensions) between latent dynamics in different session pairs after latent dynamics transformation plotted against the time intervals, for simulated data with electrode drift at 0% and 30%, from one simulation in one representative mouse with frequent stimulation. **e,** Comparing averaged correlations pooled from all session pairs for simulated data with electrode drift at 0% and 30% (*P* < 0.0001; two-tailed Wilcoxon matched-pairs signed rank test), for the mouse in **d**. **f-g,** same as in **d**-**e** but for one mouse with infrequent stimulation. In **g**, *P* < 0.0001 (two-tailed Wilcoxon matched-pairs signed rank test). **h**, Accuracies of visual information decoding from neural activities in different sessions with one single trained SVM decoder for mice with frequent visual stimulation. The inputting data are high-dimensional individual neuron level data (neuron level), latent dynamics before translation (pre-translation), and latent dynamics after transformation (transformed). **i**. Comparisons of decoding accuracies at different stages in **h** (*P* < 0.0001 for transformed vs neuron level; *P* = 0.0001 for transformed vs pre-translation; repeated measure one-way ANOVA with Geisser-Greenhouse correction and Dunnett’s multiple comparison test; n = 21 pairs). **j-k**, Confusion matrices of time window predictions from neuron level activities (**j**) and latent dynamics after transformation (**k**), for mice with frequent stimulations. **l-o,** Same as in **h-k**, but for LSTM and for infrequent mice (in **m**, *P* = 0.6997 for transformed vs neuron level; *P* = 0.4184 for transformed vs pre-translation; repeated measure one-way ANOVA with Geisser-Greenhouse correction and Dunnett’s multiple comparison test; n = 14 pairs). **p-s,** Same as in **l-o**, but for SVM (in **q**, *P* > 0.9999 for both comparisons; Friedman test and Dunn’s multiple comparison test; n = 14 pairs).

